# Studying the dynamics of the gut microbiota using metabolically stable isotopic labeling and metaproteomics

**DOI:** 10.1101/2020.03.09.982884

**Authors:** Patrick Smyth, Xu Zhang, Zhibin Ning, Janice Mayne, Jasmine I Moore, Krystal Walker, Mathieu Lavallée-Adam, Daniel Figeys

## Abstract

**Background:** The gut microbiome and its metabolic processes are dynamic systems. Surprisingly, our understanding of gut microbiome dynamics is limited. Here we report a metaproteomic workflow that involves protein stable isotope probing (protein-SIP) and identification and quantification of partially labeled peptides. We also developed a package, which we call MetaProfiler, that corrects for false identifications and performs phylogenetic and time series analysis for the study of gut microbiome functional dynamics.

**Results:** From the stool sample of five mice that were fed with ^15^N hydrolysate from *Ralstonia eutropha*, we identified 15,297 non-redundant unlabeled peptides of which 10,839 of their heavy counterparts were quantified. These peptides revealed incorporation profiles over time that were different between and within taxa, as well as between and within clusters of orthologous groups (COGs).

**Conclusions:** Our study helps unravel the complex dynamics of protein synthesis and bacterial dynamics in the mouse microbiome.

**Availability:** MetaProfiler and the bioinformatic pipeline are available at https://github.com/psmyth94/MetaProfiler.git.

## Background

Changes in the gut microbiome have been associated with a growing list of diseases including obesity[1], type II diabetes[2], inflammatory bowel disease (IBD)[3], cancer[4], neurological disorders[5] and cardiovascular issues[6]. Increasing evidences have linked the production of specific metabolites by the microbiome to these diseases[7]. The gut microbiome and its metabolic processes are not static but rather dynamic systems that continuously respond to different environmental and host derived stimuli. Surprisingly, our understanding of gut microbiome dynamics is limited. This is primarily due to the lack of analytical and bioinformatics tools to study gut microbiome dynamics. Moreover, most of the studies on gut microbiome dynamics have focused on compositional analyses of the microbiome using metagenomics[8, 9]. In contrast, metaproteomics can be used to look at functional and compositional changes in the microbiome[10].

Protein-based stable isotope probing (protein-SIP) is a well-established technique to study dynamic changes in a proteome[11]. When using protein-SIP in animals, the latter are fed food enriched with stable isotopes and their incorporation into proteins are then measured by mass spectrometry. Recently, Oberbach *et al*. reported the first study of rat gut microbiome dynamics, in which they monitored 303 bacterial peptides over a 72 hour period following the introduction of ^15^N-labelled normal chow diet (NCD) and high fat diet (HFD)[12].

Here we report a metaproteomic workflow that involved protein-SIP, identification, and quantification of partially labeled peptides, followed by computational analysis. Briefly, five 11-12 week old mice were fed ^15^N labeled hydrolysate from *Ralstonia eutropha* as mouse chow for 43 days. Stool samples from these mice were collected over 10 time points and their microbiome was analyzed by metaproteomics. Common tools available for labeled peptide indentification and quantification, such as MaxQuant[13], Census[14], and pQuant[15], are not suitable for protein-SIP experiments as they require defined mass shifts so that appropriate peptide candidates can be selected for matching against the observed MS/MS spectrum. However, there are a few tools available, such as the OpenMS tool MetaProSIP[16] and the R package ProteinTurnover[17], that can use peptide identification from light (i.e. unlabeled) peptides to identify and quantify their heavy counterparts. Thus, in order to increase the likelihood of identifying the labelled peptides over time, we spiked the unlabeled sample (time = 0) in every labeled sample to ensure the continuous presence of a light peptide, which greatly increases the ease of heavy peptide identification. We then developed a bioinformatics workflow that addresses partial ^15^N labeling in complex microbiome samples. Using this approach, 15,297 distinct peptides were identified of which 8,007 were quantified in the labelled gut microbial samples. The profile of ^15^N incorporation and the rate of newly synthesized labeled proteins over time was different between taxa of the same phylogenetic rank with the mice quickly producing proteins with ^15^N. After 43 days, the mice and microbiota had varied levels of peptides with ^15^N. The relative heavy peptide abundance at day 43 was different between clusters of orthologous groups (COG) across and within taxa. Moreover, the phylum Verrucomicrobia and the genera, Akkermansia, Lactobacillus, and Ruminococcus had not reached a plateau of protein synthesis even after 43 days. Our study revealed a complex dynamic of protein synthesis and bacterial dynamics in the mouse microbiome.

## Methods

### Mouse experiment and stool sample collection

The animal experiments were performed at the Ottawa Hospital Research Institute. The animal use protocol was approved by the Animal Care Committee at the University of Ottawa and conducted in strict accordance with the guidelines on the Care and Use of Experimental Animals of Canadian Council on Animal Care (CCAC). A total of five male C57BL/6J mice (Charles River, Sherbrooke, QC) were housed individually in the same room at 25 °C with a strict 12-h light/dark cycle. Food and water were available *ad libitum*. Mice were acclimatized to the facility for 2 weeks and fed a normal chow diet (containing 18 % fat by energy; Harlan Laboratories, Inc., Madison, WI). The diet was then switched to ^15^N-labelled SILAM Mouse Diet (Product no.: 231304650; Silantes GmbH, Munich, Germany) for 43 days, which is a ^15^N labeled hydrolysate from *Ralstonia eutropha*. This diet is a mixture of protein-free mouse chow with biomass of ^15^N-labeled *Ralstonia eutropha*. Stool samples were collected at days 0, 1, 2, 4, 8, 12, 19, 29, 34 and 43 and stored at −80 °C until analysis.

### Mouse gut microbial protein extraction and trypsin digestion

Approximately 1g of mouse stool sample were suspended in 1.5 mL of ice-cold phosphate-buffered saline (PBS, pH 7.6) with thorough vortexing (3 to 5 glass beads were added to facilitate suspension of stool pellets). The fecal slurries were centrifuged at 300g, 4 °C for 5 min. Supernatants were carefully collected, and the pellets were subjected to the above procedure three times. The supernatant for each sample was combined and then followed by three more centrifugations at 300g in 4 °C for 5 min to remove debris. The supernatant was then centrifuged at 14,000g in 4 °C for 20 min to pellet bacterial cells. The pellet was washed three times by fully re-suspending them in fresh ice-cold PBS. After the last wash, the supernatant was removed and microbial cells were lysed with 4 % (w/v) sodium dodecyl sulfate (SDS) and 6 M urea in 50 mM Tris-HCl buffer (pH 8.0). A tablet of Roche cOmplete ^™^ mini protease inhibitor tablet was added per 10ml lysis buffer. To promote microbial cell lysis, each sample was subjected to three ultrasonications (30 s each with 1 min interval on ice) using Q125 Sonicator (Qsonica, LLC) with an amplitude of 25%. Cell debris were then removed through centrifugation at 16,000g in 4 °C for 10 min. Proteins were precipitated using a 10-fold volume of acidified acetone/ethanol buffer at −20 °C overnight and washed three times using cold acetone. The precipitated proteins were then dissolved in 6 M urea/50 mM ammonium bicarbonate buffer (pH 8) for protein quantitation using DC^™^ protein assay (Bio-Rad Laboratories) and trypsin digestion.

Aliquots from each Day 0 samples were combined to generate the unlabeled sample. An equal amount of this light sample was spiked into each heavy labeled sample, generating a 1:1 ratio of light and heavy proteins for in-solution trypsin digestion. Trypsin digestion was conducted as described previously[18]. Briefly, 50 *µ*g of proteins were reduced and alkylated with 10 mM dithiothreitol (DTT) and 20 mM iodoacetamide (IAA), respectively. One microgram of trypsin (Worthington Biochemical Corp., Lakewood, NJ) was added to each sample for trypsin digestion at 37 °C overnight with agitation. The tryptic digest was desalted with a 10-*µ*m C18 column and the tryptic peptides were then eluted with 80% (v/v) acetonitrile/0.1% (v/v) formic acid. The eluted peptides were then evaporated using Savant SpeedVac^®^ Concentrator and stored at −20°C for further analysis.

### Mass Spectrometry Analysis

The dried tryptic peptides were dissolved in 0.1% (v/v) formic acid and the peptides equivalent to 2-4 *µ*g of proteins were loaded for mass spectrometry analysis on a Orbitrap Elite mass spectrometer (ThermoFisher Scientific Inc.). The separation of peptides was performed on an analytical column (75 *µ*m × 15 cm) packed with reverse phase beads (1.9 *µ*m; 120-Å pore size; Dr. Maisch GmbH, Ammerbuch, Germany). A 2-hr gradient was performed from 5 to 35 % (v/v) acetonitrile containing 0.1 % (v/v) formic acid at a flow rate of 200 nL /min. The instrument method consisted of one full MS scan from 300 to 1800 m/z using an Orbitrap mass analyzer followed by data-dependent MS/MS scan of the 20 most intense ions using a LTQ Velos Pro mass analyzer. A dynamic exclusion repeat count of 1 and repeat exclusion duration of 30 s was applied. All data were recorded with the Xcalibur package and exported as RAW format for further analysis. An overview of the wet lab portion of the workflow is shown in Figure 1.

**Figure 1.**
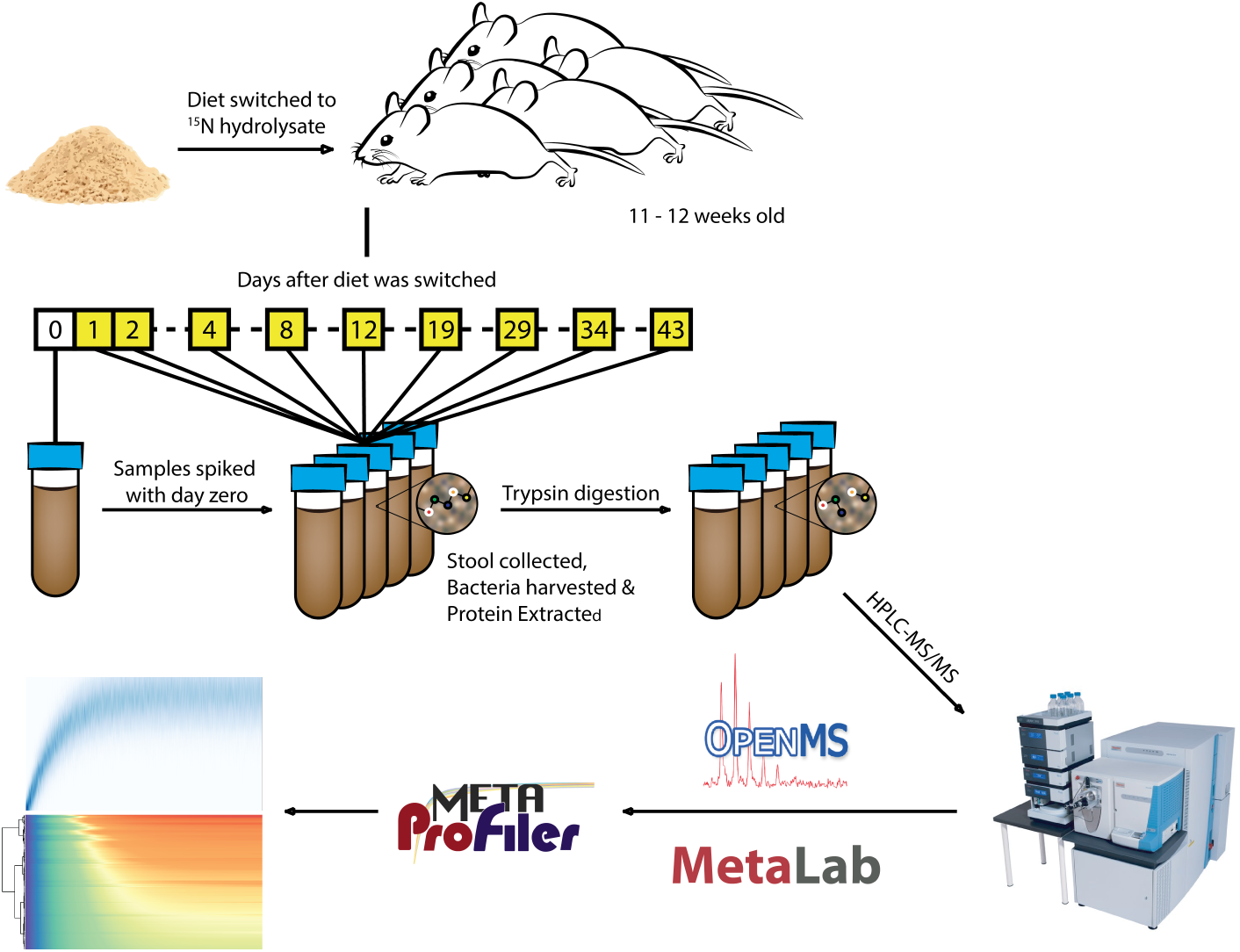
Overview of Experimental Workflow. Five 11-12 weeks old male mice were fed with a 15N labeled mouse diet, where the hydrolysate of the chemolithoautotrophic bacteria, *Ralstonia eutropha*, was the source of heavy nitrogen. Stool samples were collected at 10 different time points, were spiked with day 0 microbiome, and were processed by mass spectrometry. The mass spectrometric data was analyzed by OpenMS, MetaLab, and MetaProfiler to identify and quantify peptides and proteins and to establish profiles of ^15^N incorporations over time.

### Light and Heavy Peptide Identification and Quantification

For light peptide identification, we identified peptides and assigned them with taxonomic and functional information using MetaLab. The platform uses the MaxQuant[13] package and a target-decoy approach for peptide identification. The protein database used was derived from the catalog of the mouse gut metagenome, by Xiao *et al* [19]. A sample specific database was generated using the MS/MS clustering approach from MetaLab[19]. This database was *in silico* digested using trypsin, with two allowed missed cleavages and peptide length of 7 to 42 amino acids long. Fixed modification included carbodimethylation of cystein and variable modifications included acetylation (Protein N-term) and oxidation (M). The ppm tolerance was set at 10.

For heavy peptide identification and quantification, we used a modified template pipeline (original template available at https://sourceforge.net/projects/open-ms/files/Papers/MetaProSIP) on the KNIME platform[20], which uses tools from the open source package suite, OpenMS[21]. The difference of our pipeline from the original is that no mass calibration was used and retention time (RT) alignment used MapAlignerIdentification instead of MapAlignerPoseClustering since the light protein spike-in can be used as a point of alignment for each run. In addition, parameters were tuned so that it would be more suited for ^15^N labeling experiments. The KNIME file describing the pipeline is available at https://github.com/psmyth94/MetaProfiler.git.

Since the MaxQuant files generated from MetaLab are not compatible with the input files for the OpenMS tools, we developed an R script that can convert result files from MaxQuant into idXML. Feature selection, i.e. the group of peaks along the m/z and RT dimension that belongs to a single peptide, is performed using FeatureFinderMultiplex[22], which specializes in finding unlabeled peptide features in labeling experiments. The features were then assigned with a peptide sequence using IDMapper[22].

In order to combine all the features from each sample into a single master table, RT alignment was performed using MapAlignerIdentification[22]. Features were then linked using FeatureLinkerUnlabeledQT[23]. Linked features that contain conflicting peptide sequence information were resolved by keeping the sequence with the best score, which in this case is the peptide with the lowest Posterior Error Probability (PEP)[13] as reported by MaxQuant. This step was done with IDConflictResolver[22]. These light features were then used to identify their heavy counterparts using the MetaProSIP tool[16]. The correlation threshold was set at 0.2 as it ensured that the heavy peptide feature was selected from each mass spectrum. False discoveries were filtered from data using local false discovery rate. The overall pipeline is summarized in Figure 2A, with a more detailed pipeline available in Figure S1.

**Figure 2.**
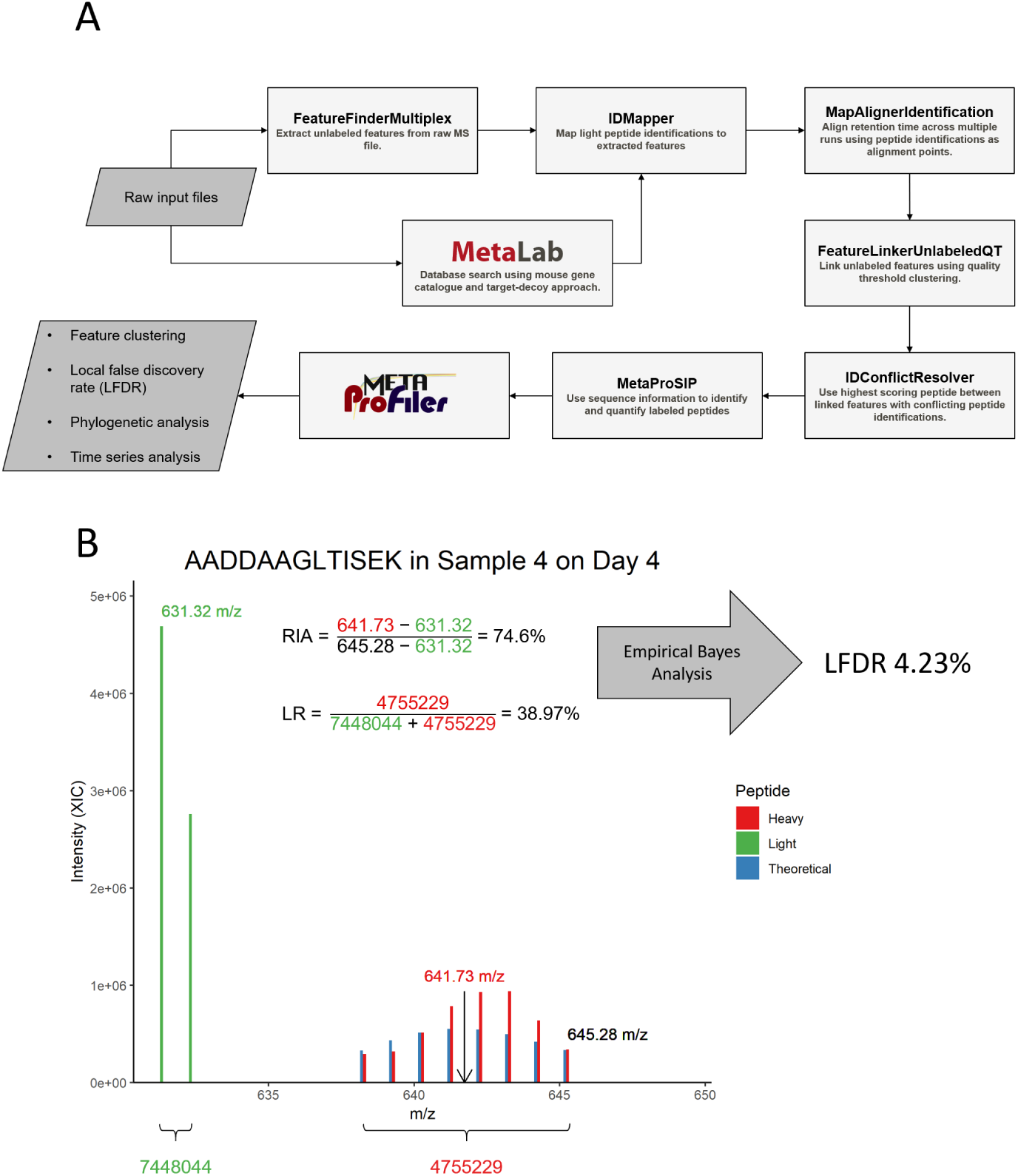
Bioinformatics Pipeline for Calculating RIA and Labeling Ratio. A) Workflow for extracting light and heavy peptide features from the raw MS files. B) Example of how RIA and LR were calculated. Once the light peptide feature was extracted from the raw MS file, a theoretical isotopic distribution (blue lines) was determined and compared against the observed peaks to extract the heavy features. Once a match was found, the m/z value from the center of the theoretical distribution was chosen for RIA calculation. LR was simply the sum of the peaks after the forth m/z peak divided by the total sum of the light and heavy features.

Quantification was done through a relative ratio between the light peptide and the estimated intensity of the heavy peptide, which is termed labeling ratio (LR) by MetaProSIP. In this case, LR measures the proportion of proteins that are produced using the heavy nitrogen from hydrolysate relative to day 0. By taking this measure over time, the rate of newly synthesized proteins that incorporate the heavy nitrogen from the hydrolysate can be estimated. Labeling ratio is calculated using equation (1)

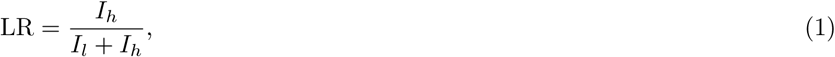

where *I*_*l*_ is the sum of the intensities of the first 4 peaks of the spectra and *I*_*h*_ is the sum of the intensities from the 5^*th*^ peak up to the total number of nitrogen in the peptide.

The elemental flux of nitrogen is measured using the average proportion of ^15^N incorporated in a peptide of interest, termed relative isotopic abundance (RIA) by MetaProSIP. By characterizing the functional and taxonomic origin of the peptide, it gives insight on how and where the stable isotopic substrate is being converted into biomass. Thus, measuring RIA can predict where this substrate is limited. RIA is calculated using equation (2)

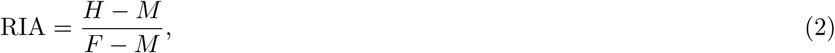

where *H* is the m/z position at the center of the theoretical isotopic distribution of the heavy peptide, *M* is the monoisotopic peak of the light peptide, and *F* is the m/z position of the fully labeled peptide. An example of how RIA and LR is measured is illustrated in Figure 2B.

### Filtering Heavy Peptide Features using Local False Discovery Rate

One particular issue with doing biological interpretation from proteomic data is that some peptide identifications can be the result of a false positive (in our case, heavy features quantified from co-eluting peptides or noisy peaks). Thus, confidence scores are often reported with proteomic data so that confident peptide identifications can be obtained. In order to estimate a confidence score for the heavy peptide feautres obtained, we can use the idea that the probability of obtaining high RIAs or LRs at early timepoints is lower than the probabiltiy of obtaining high RIAs or LRs at later timepoints. In addtion, we can also use the idea that heavy peptide features from false discoveries would (usually) have equal probability of obtaining an RIA or LR at any given timepoint. With this in mind, a confidence score can be obtained by using an empirical Bayesian approach to calculate a local false discovery rate (LFDR). LFDR, formulated by Efron *et al* [24], is defined as the probability of the hypothesis being null given the observed data and it is calculated using three parameters: i) the probability of getting a false discovery, ii) the density of the data for false discoveries, and iii) the mixture density of the data for false and true discoveries (see supplemental information for full equation). In our study, the density of RIA or LR for false discoveries was empirically estimated from time zero using kernel density estimation (KDE) since all of the heavy peptide features quantified at time zero are false. Similarly, the probability of false discoveries at a given timepoint can be estimated using the total number of heavy features identified at time zero and the total number of heavy features identified at a particular timepoint. Finally, the mixture density of RIA or LR for false and true discoveries was measured at each timepoint using KDE. In order to calculate the joint *a posteriori* probability of false discovery given that both RIA and LR is observed, we assume that both random variables are conditionally independent given the hypothesis so that the chain rule can be applied. Once calculated, a threshold can then be set on the LFDR to obtain confident heavy peptide features. For this study, features with less than 10% LFDR were used in downstream analysis. It must be noted that the disadvantage to using this approach is that heavy peptide features quantified from unlikely events−for example, those that have both high RIA and LR at early timepoints−will often have high LFDR. Thus, these features will be lost in those timepoints when setting the LFDR threshold low.

### Combining Pseudo-Features

MetaProSIP has the option to use unassigned peptides, which are peptides identified by an MS/MS spectrum but not assigned to a feature, by forming them into pseudo-features. These pseudo-features are then associated with heavy peptides by MetaProSIP using sequence information. Since several MS/MS spectra can originate from the same peptide feature, we built a tool written in C++ and integrated into R using Rcpp[25] that performs quality threshold clustering to merge these redundant features. In this study, all features that were within 1% RIA, 30s RT, and 1% LR away from their respective mean in the cluster were merged into a single feature.

### Data Imputation

In order to predict the RIA or LR at missing timepoints, we used equation derived from a three-compartment model and developed by Guan *et al* [26](see additional file 3 for full equation). The four constants that define the three-compartment model are estimated using the C++ code from RcppDE[27], which is a package that uses a differential evolutionary (DE) algorithm for minimizing a cost function. The advantage of using DE was that it struck a balance between the robustness of an exhaustive search and the speed of a gradient descent. The least squares approach was used as the cost function. This approach was evaluated using a leave-one-out cross validation (LOOCV) approach.

### Data Analysis and Visualization

Data analysis was performed using our developed tool MetaProfiler. All plots were generated using ggplot2[28] or ComplexHeatmap[29]. The phylogenetic tree was generated with GraPhlAn[30]. The optimal number of clusters from the hierarchical clustering approach was determined by the R package, NbClust[31], which reports the best number of clusters by majority vote from 30 clustering indices. To test for over-representation of taxa in each hierarchical cluster, a hypergeometric test was used to assess significance. The total number of non-redundant peptides with more than 4 timepoints were used as background for the hypergeometric test. The p-values were adjusted using benjamini-hochberg (BH) procedure. The package is available at https://github.com/psmyth94/MetaProfiler.git. The mass spectrometry proteomics data have been deposited to the ProteomeXchange with the dataset identifier PXD017451.

## Results and Discussion

We investigated the *in vivo* dynamics of the microbiome using metaproteomics coupled with protein-SIP. Briefly, five mice were fed ^15^N labeled hydrolysate from *Ralstonia eutropha* for 43 days, stool samples were collected over 10 time points and their microbiome was analyzed by metaproteomics (Figure 1). One challenge when dealing with ^15^N labeled proteomic data, in particular from the microbiome, is the difficulty in confidently identifying peptides from MS/MS spectra due to the partial incorporation of ^15^N[16, 17]. To address this issue, unlabeled microbiome samples (time = 0) were spiked in every partially labeled sample to ensure the presence of the corresponding light peptide and confident peptide identification. By using this experimental design, 10,839 non-redundant heavy peptides corresponding to 2,075,033 features were quantified in the labelled gut microbial samples from the 15,297 non-redundant light peptide identifications by pipeline. The pipeline for extracting the heavy peptide features is detailed in the Method section as well as summarized in Figure 2A. Peptides with at least two RIAs, one with a value below one over the maximum number of nitrogen, 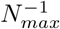, which corresponds to the light peptide, as well as an RIA above *N*^−1^, which corresponds to the heavy peptide. After filtering at a threshold level of 10% LFDR, this resulted in 8,007 heavy labeled peptides corresponding to 493,165 features across all samples (Table S1).The reason for clustering the features together is because this process significantly decreases the number of data and facilitates downstream analysis. In addition, since the median LFDR for day 1, 2, 4, 8, 12, 19, 29, 34, and 43 are 19%, 11%, 9%, 8%, 8%, 9%, 9%, 9%, and 8%, respectively, the threshold was set at 10% so that earlier timepoints would not be filtered out completely. The reason for LFDR being higher in earlier timepoints is because low RIAs are more likely to be false as MetaProSIP may sometimes mistake the light peptide feature as being overlapped with the heavy peptide feature at time zero. This is why a higher density of RIA is observed for false discoveries in earlier timepoints in Figure S2A. In addition, quantified features that are the result of noise will often be low in LR, as it is seen in Figure S2B.

### Taxonomic and functional characterization of the heavy labeled peptides in mouse gut microbiome

To characterize the functions and taxonomic origin of heavy labeled proteins in microbiome samples, we used the confidently quantified heavy peptides at day 29 to 43. These days were chosen by determining which day the LR starts to plateau. Plateau detection was done using a sliding window method, where a window of three timepoints moves over the LR of each peptide and then computes the standard deviation of the LR within the window. The average standard deviation across all peptides at each permutation are 10.51% (day: 1, 2, 4), 9.78% (day: 2, 4, 8), 8.87% (day: 4, 8, 12), 8.55% (day: 8, 12, 19), 7.88% (day: 12, 19, 29), 7.48% (day: 19, 29, 34), and 6.9% (day: 29, 34, 43). A two sampled t-test showed that the standard deviation in the last window with day 29, 34, and 43 is significantly less than all other windows (p-value = 2×10^−106^, 2×10^−73^, 1×10^−34^, 2×10^−23^, 1×10^−8^, and 0.0008, respectively; BH procedure). Thus, the LRs at these timepoint reflect the maximum proportion of active proteins that the taxon/function can attain in the mouse gut. An LR higher than 50% denotes higher production of this protein compared to day 0. An LR lower than 50% is either reflection of a decreased production of that protein or a reflection that nitrogen is obtained through an alternative source other than the hydrolysate. The distribution of RIA and LR over time is show in Figure S3A and B, respectively.

We first mapped the taxonomic LR to a phylogenetic tree using GraPhlAn (Figure 3). Taxonomic LR was taken as the average across all its distinct peptides. Proteins from mice (Chordata) were within the highest peptide LR ratios (0.572 ± 0.015 at 95% CI). This is not particularly surprising as dietary proteins are the most common source of nitrogen for mammals. Similarly, Verrucomicrobia, a group of mucin degraders who rely on the host for nutrients, had a similar LR level (0.555 ± 0.067 at 95% CI). Three of the four dominant phyla in the gut have mid-range LR levels including Proteobacteria (0.401 ± 0.047 at 95% CI), Bacteroidetes (0.434 ± 0.009 at 95% CI), and Actinobacteria (0.341 ± 0.140 at 95% CI). Interestingly, the forth and most dominant phyla, Firmicutes, had significantly lower LR (0.335 ± 0.006 at 95% CI), than the host cells (p-value=9×10^−51^; two sample t-test; BH procedure) and Bacteroidetes (p-value=4×10^−25^; two sample t-test; BH procedure). A possible explanation is that they do not process the hydrolysate efficiently and may obtain their nitrogen source elsewhere, such as from fiber[32] or metabolites from other microorganism in the microbial community. While most of the taxa belonging to Firmicutes had low LR, the LR of the lactic acid bacteria group (e.g. Lactobacillus; 0.535 ± 0.057 at 95% CI) was markedly higher, suggesting potential differences in nutritional mechanisms of lactic acid bacteria with other Firmicutes species. The mean LR of each taxon are available in Table S2. To further examine the functional distributions of ^15^N labeled proteins, taxa were assigned to categories of COGs (Figure 4) using the leading razor proteins associated with their distinct peptides, as reported from MaxQuant. Functions of “carbohydrate transport and metabolism” and “amino acid transport and metabolism”, “energy production and conversion”, as well as “translation, ribosomal structure, and biogenesis” were among the categories with the highest number of heavy peptide associated to them. The composition of functions was similar to previous metaproteomic studies on the gut microbiome[33], indicating that the labeling might not be biased to specific functional categories. The mean LR of each taxon and their corresponding COG category are available in Table S3 and the list of protein names are available in Table S4. The LR of proteins belonging to the functional group “amino acid transport and metabolism” corresponded well with the overall LR of heavy peptides in each taxon (Pearson correlation = 0.80). This makes sense as those which efficiently degrade the hydrolysate will generally see a higher level of heavy peptides. However, there were a few taxa which displayed high overall levels of LR without having peptides belonging to this category. In particular, *Akkermansia muciniphila*, showed high LR in the functional group “carbohydrate transport and metabolism” (0.558 ± 0.108 at 95% CI), which was consistent with its mucin degrading ability. Another taxon which showed this pattern is Lactobacillus (0.595 ± 0.048 at 95% CI). Although not as well characterized, several species of this lineage have been shown to possess proteins that degrade mucin[34]. In addition, both Akkermansia and Lactobacillus were high in “translation, ribosomal structure and biogenesis” (0.667 ± 0.081 at 95% CI) and 0.462 ± 0.150 at 95% CI), respectively), which indicates that both are rapid growers.

**Figure 3.**
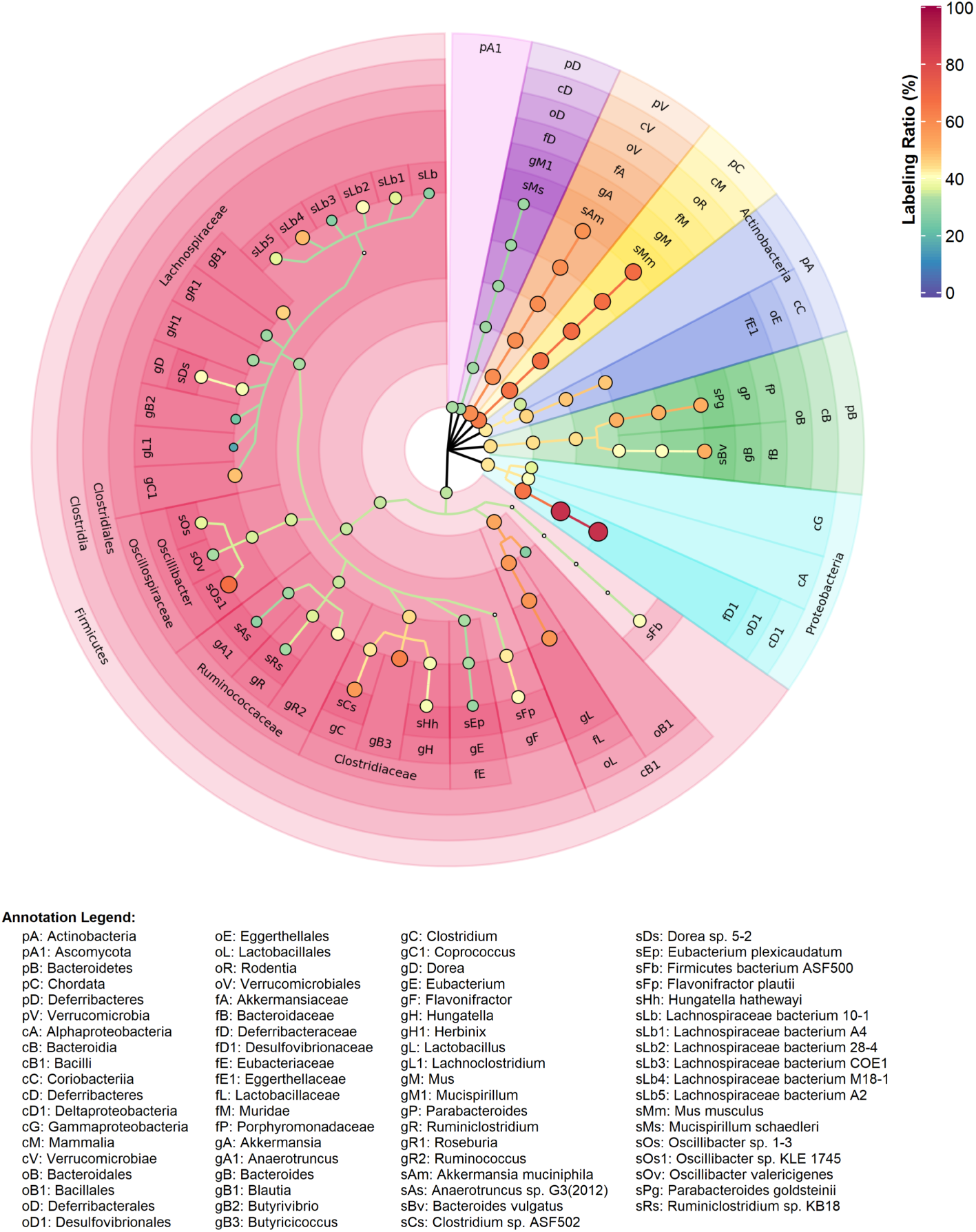
Labeling Ratio of Proteins in Mouse Microbiome on Day 43. Phylogenetic tree where size and color of the nodes relates to the LR of the taxon; color of the cells corresponds to the phylum lineage; and the transparency denotes the rank. For nodes with annotations, the key follows the convention: [first letter of phylogenetic rank][initials][unique identifier]. Nodes are reported when the number of distinct peptides is greater than 3 and the adjusted p-value from a one-sample t-test is below 0.05.

**Figure 4.**
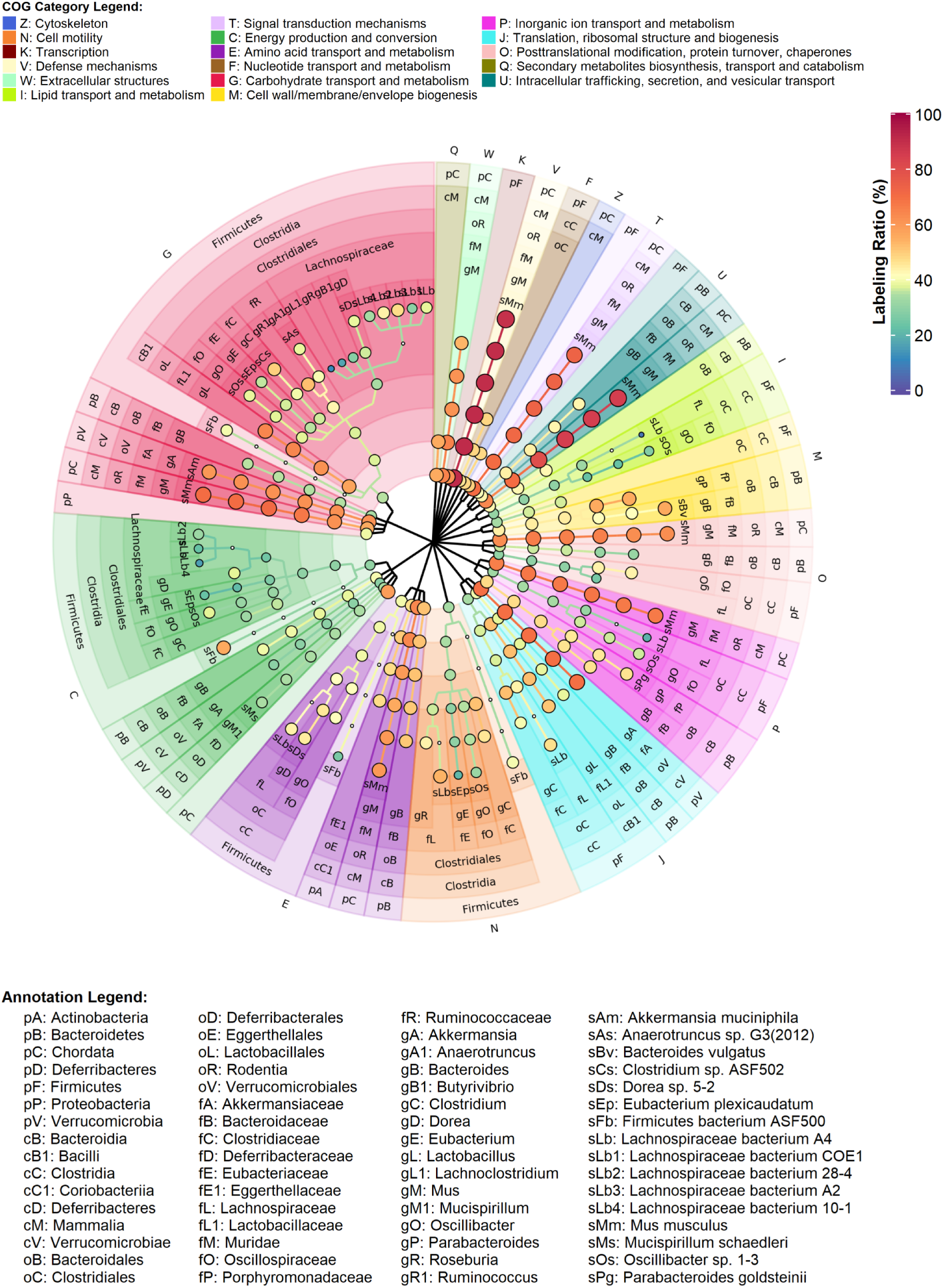
Functional Distribution of Heavy Labeled Proteins in Mouse Gut Microbiome on Day 43. The tree describes the functional distribution of each taxon. The size and color of the nodes relates to the LR of the taxon; color of the cells corresponds to the functional COG category; and the transparency denotes the rank. Nodes were annotated and reported in a similar fashion to figure 3.

The host showed high levels of heavy peptides in most categories. In particular, peptides from “intracellular trafficking, secretion, and vesicular transport” (0.610 ± 0.045 at 95% CI), which mostly includes calcium-dependent phospholipid-binding proteins, are among the highest ^15^N incorporators. On the other hands, peptides from “secondary metabolites biosynthesis, transport, and catabolism” (0.436 ± 0.209 at 95% CI), which mostly includes multidrug resistant proteins, were among the lowest incorporators of ^15^N. This was reasonable as these proteins are only expressed in response to drugs[35, 36].

### Dynamics of ^15^N Incorporation into Peptides of the Mouse Gut Microbiota

We next explored the ^15^N incorporation profiles over time. In particular, we focused on the subset of 3,420 peptides that had sufficient data points to perform time-series modeling (observed in at least 5 time points and in at least 3 samples). We assessed whether the datasets could be modeled using a three-exponential regression model, previously formulated by Guan et al. for computing protein turnover[26]. This regression model predicted missing values with an average RMSE of 9% for LR and 6% for RIA. The results before and after imputation is available in Figure S5.

We then evaluated the overall nitrogen flux in mice gut microbiome (Figure 5A and Table S5). Nitrogen incorporation appeared to be rapid in the first few days but then slowed down (Figure S4A). Additionally, the vast majority of peptides did not reach an RIA of 100%, where the interquartile range was between 88.9% and 91.8% at day 43. Nitrogen fixation could provide another source of unlabeled nitrogen; however, very few mammalian gut microbes are capable of nitrogen fixation[37, 38]. It is more likely that the microbiota cells obtain light amino acids via host protein degradation, and *de novo* synthesis[17]. Overtime, the nitrogen pool will gradually grow towards heavy isotopes as the host increasingly incorporates the labeled heavy nitrogen. To further examine whether there were differences in nitrogen incorporation rate of different peptides, we performed clustering using the first four time points (day 1, 2, 4, and 8), where higher RIA variations were observed. Two distinct clusters (RIAC1, RIAC2) were identified with the peptide in cluster RIAC3 showed the fastest isotope incorporation. Interestingly, a noticeable lag in incorporation was present in cluster RIAC1 from day 1 to day 2. Hypergeometric tests showed that *Parabacteroides goldsteinii* and host cells are significantly over-represented in cluster RIAC1 (p-value = 4×10^−7^ and 8×10^−3^, respectively; BH procedure; Table S7). To investigate this further, a pairwise t-test between taxa using their distinct heavy peptide RIA accompanied by BH procedure was performed to identify which species had significantly less nitrogen incorporation at day 2. Consistent with the hypergeometric tests, mouse cells are significantly less than *Eubacterium plexicaudatum* (p-value=1×10^−2^), *Firmicutes bacterium ASF500* (p-value=1×10^−2^), *Lachnospiraceae bacterium 10-1* (p-value=4×10^−2^), while *Parabacteroides goldsteinii* is significantly less than *Akkermansia muciniphila* (p-value=4×10^−2^), *Clostridium sp. ASF502* (p-value=4×10^−2^), *Eubacterium plexicaudatum* (p-value=9×10^−5^), *Firmicutes bacterium ASF500* (p-value=9×10^−5^), *Lachnospiraceae bacterium 10-1* (p-value=8×10^−4^), *Lachnospiraceae bacterium 28-4* (p-value=3×10^−3^), *Lachnospiraceae bacterium A4* (p-value=1×10^−2^), *Mus musculus* (p-value=1×10^−2^), *Oscillibacter sp. 1-3* (p-value=2×10^−4^). One possible explanation for this observation was that heavy nitrogen availability was delayed to certain strains of *Parabacteroides goldsteinii* in the gut. However, once heavy nitrogen became readily available to them, these strains started to catch up with the earlier incorporators.

**Figure 5.**
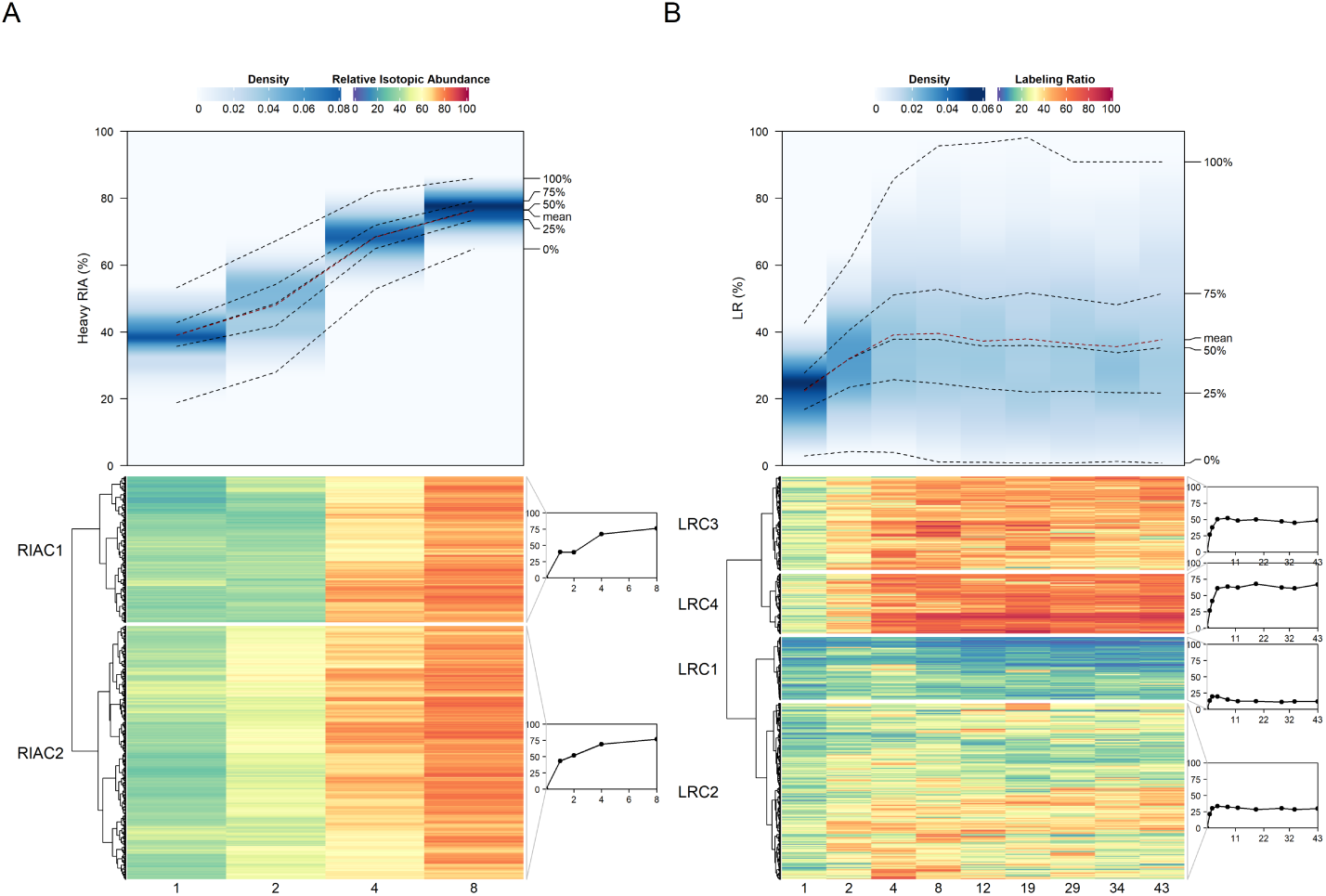
Isotope Incorporation Profile. A) The top graph shows the distribution of RIA at each time point. Dark blue denotes a high density and light blue denotes a low density. At the bottom is a heatmap where the rows are peptides and the columns are the days at which the RIA was recorded. B) Similar to A except that it denotes labeling ratio over time. Both heatmaps were clustered using Ward’s minimum variance, as implemented in [39], and Euclidean distance.

In order to group the different rates of newly synthesized proteins, hierarchical clustering of the LR over time was performed (Figure 5B and Table S6). A total of four clusters were identified (LRC1, LRC2, LRC3, LRC4). Peptides from cluster LRC1 showed the lowest LR across all time points. Strikingly, those in cluster LRC1 showed a characteristic decline after day 4. Since the heavy nitrogen was provided by the hydrolysate, this indicates that the source of nitrogen for these taxa might differ. Hypergeometric tests showed that Firmicutes is significantly over-represented in this cluster (p-value = 4×10^−5^; BH procedure; Table S7). Cluster LRC2, LRC3, and LRC4, on the other hand, remained stable until day 43 after the 4^*th*^ day of ^15^N diet feeding, indicating an equilibrium state was reached by these peptides. Peptides from cluster LRC4 reached the highest LR ratio (0.673 ± 0.009 at 95% CI). In addition, the hypergeometric tests showed that the taxa over-represented in cluster LRC4 are mouse cells (p-value = 4×10^−36^; BH procedure) and *Bacteroides vulgatus* (p-value = 0.01; BH procedure) and in LRC3 are *Parabacteroides goldsteinii* (p-value = 2×10^−6^; BH procedure). This is consistent with Figure 3, where the majority of the mouse cells had high LR. Altogether, these findings suggest that the rates of newly synthesized proteins varied among different taxa.

### ^15^N incorporation rate differed according to microbial phylogeny

To further investigate whether the incorporation rate of nitrogen differed between microorganisms in mouse gut microbiome, we mapped all the above peptides to taxa and identified 3 superkingdom, 2 kingdom, 9 phylum, 12 class, 12 order, 15 family, 17 genus, and 21 species that have more than 3 distinctive peptides. RIA and LR of each taxon at each time point were then calculated using the average of all distinctive peptides (see additional file 1 for average RIA and LR over time for all taxa). At the phylum level, the RIA profiles over time were similar among all the identified seven phyla (Figure 6A), with the exception of host cells (Chordata) and Bacteroidetes between day 1 and 2. This further supports the hypergeometric tests performed in the clusters presented in Figure 5A.

**Figure 6.**
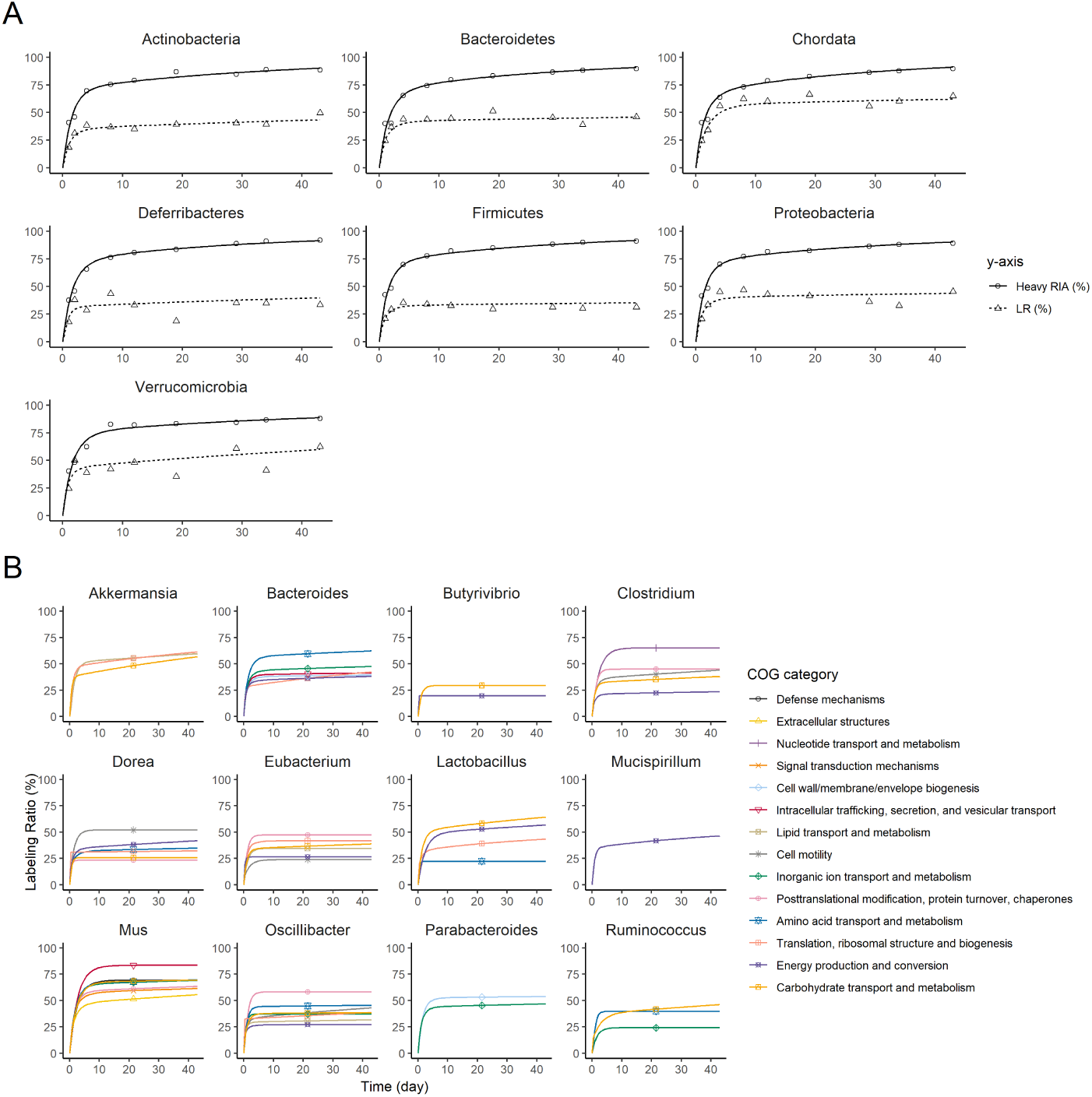
Taxon-Specific RIA and LR Profiles Over Time. (A) phylum level RIA and LR profiles over time; (B) Average COG LR profiles over time of the 12 most abundant genera.

Obvious different LR profiles were also observed in Figure 6A. The synthesis rate for Firmicutes clearly exhibited a similar pattern found in cluster LRC2 in Figure 5B, and reached a lower plateau than other abundant phyla, such as Bacteroidetes, Proteobacteria and the mice host. In agreement with the results obtained above, the mice proteins showed the highest LR after 4 days of ^15^N diet. These findings suggested that the incorporation of ^15^N to the host proteins was faster than the microbiome. To investigate this further, we profiled the LR over time of the most abundant genera and profiled the average LR overtime across all their COG categories. The pattern that we observed in Figure 6B in COG category “amino acid transport and metabolism” for mice (Mus) was consistent with amino acids as a common source of nitrogen for animal cells.

The LR curve of Verrucomicrobia showed a characteristic increasing trend and never fully reached a plateau over 43 days. This is also observed in the genus, Akkermansia (Figure 6B), for category “translation, ribosomal structure and biogenesis” and “carbohydrate transport and metabolism”. A similar pattern was observed for Lactobacillus in these categories as well, which further provided support of their mucin degrading capabilities. Interestingly, Ruminococcus also displayed this pattern in “carbohydrate transport and metabolism” and previous work have suggested that these are mucin degraders as well[40]. It is possible that these taxa may play important roles in foraging nitrogen from the host to the microbiome. Thus, our findings may suggest that these taxa, especially *A. muciniphila*, are keystone species for the metabolic flux of nitrogen in the microbiome.

## Conclusions

We investigated the *in vivo* dynamics of the microbiome using metaproteomics coupled with protein-SIP. We developed an approach that combined spike-in of samples with a new bioinformatics pipeline and a new bioinformatics tool called MetaProfiler. This bioinformatics approach greatly enhanced the number of peptides identified (15,297), and quantified (10,839; 8,007 with LFDR *<* 10%) with ^15^N incorporations. Our study revealed a complex dynamic of protein synthesis and bacterial dynamics in the mouse microbiome.

## Supporting information

Average Taxonomic LR and RIA Over Time

## Ethics approval and consent to participate

The animal use protocol was approved by the Animal Care Committee at the University of Ottawa and conducted in strict accordance with the guidelines on the Care and Use of Experimental Animals of Canadian Council on Animal Care (CCAC).

## Consent for publication

Not applicable.

## Availability of data and material

MetaProfiler and KNIME pipeline is available at https://github.com/psmyth94/MetaProfiler.git. The mass spectrometry data and pipeline results are available in ProteomeXchange with the dataset identifier PXD017451.

## Authors’ contributions

P. Smyth took the lead in writing the manuscript. X. Zhang, Z. Ning, J. Mayne, M. Lavallée-Adam, and D. Figeys provided critical feedback on the manuscript. MetaProfiler was written by P. Smyth, with supervision from M. Lavallée-Adam. The KNIME pipeline was developed by the OpenMS team, and modified by P. Smyth and Z. Ning. Experimental protocol was designed by X. Zhang. Mouse experiment, stool sample collection, protein extraction, trypsin digestion, and mass spec analysis were performed by X. Zhang, J. Mayne, J. I. Moore, and K. Walker.

## Acknowledgements

Not applicable.

## Additional Files

Additional file 1 — Average taxon RIA and LR over time

Average taxon RIA and LR over time. Each taxon has geater than three number of distinct peptides.

Additional file 2 — Supplemental

figures Figure S1-4.

Additional file 3 — Supplemental tables

Table S1-7.

TableS1 – Master Table

Table which includes the light and heavy relative isotopic abundance (RIA) and intensity (INT) of each peptide, along with their labeling ratio (LR), correlation score (Cor.), local false discovery rates (LFDR), proteins, functional and taxonomic annotations.

TableS2 – Average Taxonomic LR Over Time

The average LR of each taxon, including the confidence (conf.) interval and p-value from the one sample t-test.

TableS3 – Average Taxonomic and Functional LR Over Time

Similar to table S2, except the value is now the average taxonomic LR in each clusters of orthologous groups (COG) categories.

TableS4 – Taxon to Protein Names

The list of protein names identified from each taxon, including their LR value in day 29, 34, and 43.

TableS5 – RIA Clusters

The RIA at day 1, 2, 4, 8, 12, 19, 29, 34, and 43 of each peptide along with their corresponding cluster ID.

TableS6 – LR Clusters

Similar to Table S5, but with LR instead.

TableS7 – Hypergeometic Tests

The hypergeometric test results for over-representation of taxa in each hierarchical cluster. Significane was achieved when the adjusted p-value was below 0.05.

## References

1. Maruvada P, Leone V, Kaplan LM, Chang EB. The Human Microbiome and Obesity: Moving beyond Associations. Cell Host Microbe. 2017;22(5):589–599. Available from: https://www.ncbi.nlm.nih.gov/pubmed/29120742.

2. Komaroff AL. The Microbiome and Risk for Obesity and Diabetes. JAMA. 2017;317(4):355–356. Available from: https://www.ncbi.nlm.nih.gov/pubmed/28006047.

3. Franzosa EA, Sirota-Madi A, Avila-Pacheco J, Fornelos N, Haiser HJ, Reinker S, et al. Gut microbiome structure and metabolic activity in inflammatory bowel disease. Nat Microbiol. 2019;4(2):293–305. Available from: https://www.ncbi.nlm.nih.gov/pubmed/30531976.

4. Helmink BA, Khan MAW, Hermann A, Gopalakrishnan V, Wargo JA. The microbiome, cancer, and cancer therapy. Nat Med. 2019;25(3):377–388. Available from: https://www.ncbi.nlm.nih.gov/pubmed/30842679.

5. Tremlett H, Bauer KC, Appel-Cresswell S, Finlay BB, Waubant E. The gut microbiome in human neurological disease: A review. Ann Neurol. 2017;81(3):369–382. Available from: https://www.ncbi.nlm.nih.gov/pubmed/28220542.

6. Tang WH, Hazen SL. The Gut Microbiome and Its Role in Cardiovascular Diseases. Circulation. 2017;135(11):1008–1010. Available from: https://www.ncbi.nlm.nih.gov/pubmed/28289004.

7. Van Treuren W, Dodd D. Microbial Contribution to the Human Metabolome: Implications for Health and Disease. Annu Rev Pathol. 2020;15:345–369. Available from: https://www.ncbi.nlm.nih.gov/pubmed/31622559.

8. Halfvarson J, Brislawn CJ, Lamendella R, Vazquez-Baeza Y, Walters WA, Bramer LM, et al. Dynamics of the human gut microbiome in inflammatory bowel disease. Nat Microbiol. 2017;2:17004. Available from: https://www.ncbi.nlm.nih.gov/pubmed/28191884.

9. Schirmer M, Franzosa EA, Lloyd-Price J, McIver LJ, Schwager R, Poon TW, et al. Dynamics of metatranscription in the inflammatory bowel disease gut microbiome. Nat Microbiol. 2018;3(3):337–346. Available from: https://www.ncbi.nlm.nih.gov/pubmed/29311644.

10. Zhang X, Chen W, Ning Z, Mayne J, Mack D, Stintzi A, et al. Deep Metaproteomics Approach for the Study of Human Microbiomes. Anal Chem. 2017;89(17):9407–9415. Available from: https://www.ncbi.nlm.nih.gov/pubmed/28749657.

11. Jehmlich N, Vogt C, Lunsmann V, Richnow HH, von Bergen M. Protein-SIP in environmental studies. Curr Opin Biotechnol. 2016;41:26–33. Available from: https://www.ncbi.nlm.nih.gov/pubmed/27116035.

12. Oberbach A, Haange SB, Schlichting N, Heinrich M, Lehmann S, Till H, et al. Metabolic in Vivo Labeling Highlights Differences of Metabolically Active Microbes from the Mucosal Gastrointestinal Microbiome between High-Fat and Normal Chow Diet. J Proteome Res. 2017;16(4):1593–1604. Available from: https://www.ncbi.nlm.nih.gov/pubmed/28252966.

13. Cox J, Mann M. MaxQuant enables high peptide identification rates, individualized p.p.b.-range mass accuracies and proteome-wide protein quantification. Nat Biotechnol. 2008;26(12):1367–72. Available from: https://www.ncbi.nlm.nih.gov/pubmed/19029910.

14. Park SKR, Aslanian A, McClatchy DB, Han X, Shah H, Singh M, et al. Census 2: isobaric labeling data analysis. Bioinformatics. 2014;30(15):2208–2209.

15. Liu C, Song CQ, Yuan ZF, Fu Y, Chi H, Wang LH, et al. pQuant improves quantitation by keeping out interfering signals and evaluating the accuracy of calculated ratios. Analytical chemistry. 2014;86(11):5286–5294.

16. Sachsenberg T, Herbst FA, Taubert M, Kermer R, Jehmlich N, von Bergen M, et al. MetaProSIP: automated inference of stable isotope incorporation rates in proteins for functional metaproteomics. Journal of proteome research. 2015;14(2):619–627.

17. Fan KT, Rendahl AK, Chen WP, Freund DM, Gray WM, Cohen JD, et al. Proteome scale-protein turnover analysis using high resolution mass spectrometric data from stable-isotope labeled plants. Journal of proteome research. 2016;15(3):851–867.

18. Zhang X, Li L, Mayne J, Ning Z, Stintzi A, Figeys D. Assessing the impact of protein extraction methods for human gut metaproteomics. J Proteomics. 2018;180:120–127. Available from: https://www.ncbi.nlm.nih.gov/pubmed/28705725.

19. Xiao L, Feng Q, Liang S, Sonne SB, Xia Z, Qiu X, et al. A catalog of the mouse gut metagenome. Nature biotechnology. 2015;33(10):1103.

20. Berthold MR, Cebron N, Dill F, Gabriel TR, Kötter T, Meinl T, et al. KNIME-the Konstanz information miner: version 2.0 and beyond. AcM SIGKDD explorations Newsletter. 2009;11(1):26–31.

21. Röst HL, Sachsenberg T, Aiche S, Bielow C, Weisser H, Aicheler F, et al. OpenMS: a flexible open-source software platform for mass spectrometry data analysis. Nature methods. 2016;13(9):741.

22. Alka O, Sachsenberg T, Bichmann L, Pfeuffer J, Weisser H, Wein S, et al. OpenMS for open source analysis of mass spectrometric data. PeerJ Preprints. 2019;7:e27766v1.

23. Weisser H, Choudhary JS. Targeted feature detection for data-dependent shotgun proteomics. Journal of proteome research. 2017;16(8):2964–2974.

24. Efron B, Tibshirani R, Storey JD, Tusher V. Empirical Bayes analysis of a microarray experiment. Journal of the American statistical association. 2001;96(456):1151–1160.

25. Eddelbuettel D, François R, Allaire J, Ushey K, Kou Q, Russel N, et al. Rcpp: Seamless R and C++ integration. Journal of Statistical Software. 2011;40(8):1–18.

26. Guan S, Price JC, Ghaemmaghami S, Prusiner SB, Burlingame AL. Compartment modeling for mammalian protein turnover studies by stable isotope metabolic labeling. Analytical chemistry. 2012;84(9):4014–4021.

27. Eddelbuettel D. RcppDE: Global optimization by differential evolution in C++. R package version 01 0, URL http://CRANR-projectorg/package=RcppDE. 2010;.

28. Wickham H. ggplot2: elegant graphics for data analysis. Springer; 2016.

29. Gu Z, Eils R, Schlesner M. Complex heatmaps reveal patterns and correlations in multidimensional genomic data. Bioinformatics. 2016;32(18):2847–2849.

30. Asnicar F, Weingart G, Tickle TL, Huttenhower C, Segata N. Compact graphical representation of phylogenetic data and metadata with GraPhlAn. PeerJ. 2015;3:e1029.

31. Charrad M, Ghazzali N. Package ‘nbclust’. Journal of statistical software. 2014;61:1–36.

32. Deehan EC, Duar RM, Armet AM, Perez-Munoz ME, Jin M, Walter J. Modulation of the Gastrointestinal Microbiome with Nondigestible Fermentable Carbohydrates To Improve Human Health. Microbiol Spectr. 2017;5(5). Available from: https://www.ncbi.nlm.nih.gov/pubmed/28936943.

33. Maier TV, Lucio M, Lee LH, VerBerkmoes NC, Brislawn CJ, Bernhardt J, et al. Impact of Dietary Resistant Starch on the Human Gut Microbiome, Metaproteome, and Metabolome. mBio. 2017;8(5). Available from: https://www.ncbi.nlm.nih.gov/pubmed/29042495.

34. McDonald ND, Lubin JB, Chowdhury N, Boyd EF. Host-Derived Sialic Acids Are an Important Nutrient Source Required for Optimal Bacterial Fitness In Vivo. mBio. 2016;7(2):e02237–15. Available from: https://www.ncbi.nlm.nih.gov/pubmed/27073099.

35. Klaassen CD, Aleksunes LM. Xenobiotic, bile acid, and cholesterol transporters: function and regulation. Pharmacol Rev. 2010;62(1):18 – 23. Available from: https://www.ncbi.nlm.nih.gov/pubmed/20103563.

36. Mottino AD, Hoffman T, Jennes L, Vore M. Expression and localization of multidrug resistant protein mrp2 in rat small intestine. J Pharmacol Exp Ther. 2000;293(3):717–23. Available from: https://www.ncbi.nlm.nih.gov/pubmed/10869369.

37. Reese AT, Pereira FC, Schintlmeister A, Berry D, Wagner M, Hale LP, et al. Microbial nitrogen limitation in the mammalian large intestine. Nature microbiology. 2018;3(12):1441–1450.

38. Wostmann BS. The germfree animal in nutritional studies. Annual review of nutrition. 1981;1(1):257–279.

39. Murtagh F, Legendre P. Ward’s hierarchical agglomerative clustering method: which algorithms implement Ward’s criterion? Journal of classification. 2014;31(3):274–295.

40. Crost EH, Le Gall G, Laverde-Gomez JA, Mukhopadhya I, Flint HJ, Juge N. Mechanistic insights into the cross-feeding of Ruminococcus gnavus and Ruminococcus bromii on host and dietary carbohydrates. Frontiers in microbiology. 2018;9:2558.

