## Supplementary material for "Studying the dynamics of the gut microbiota using metabolically stable isotopic labeling and metaproteomics": Average Taxonomic LR and RIA Over Time

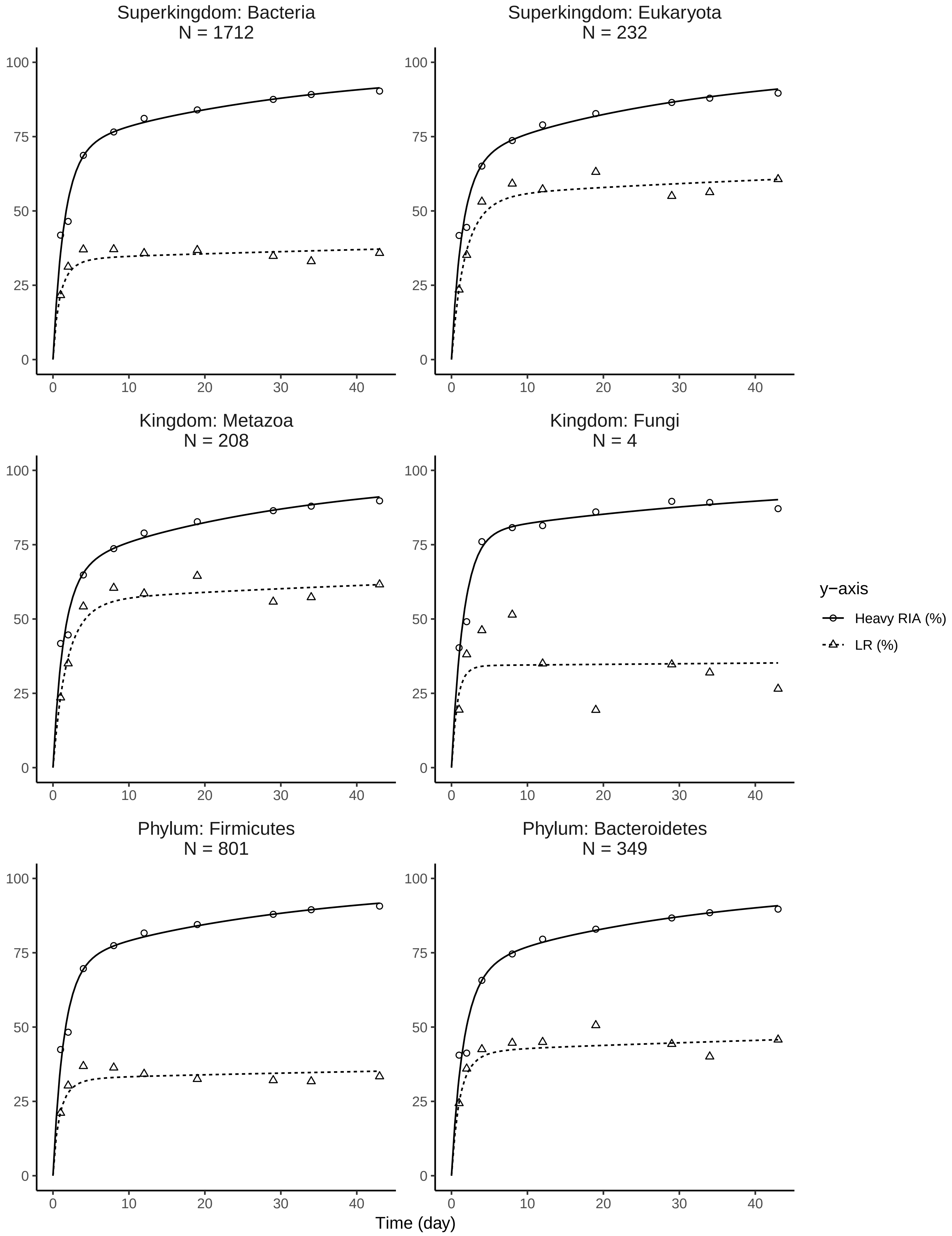
Superkingdom: Bacteria
N = 1712
Superkingdom: Eukaryota
N = 232
Kingdom: Metazoa
N = 208
Kingdom: Fungi
N = 4
Phylum: Firmicutes
N = 801
Phylum: Bacteroidetes
N = 349
y−axis
Heavy RIA (%)
LR (%)
Time (day)
0
25
50
75
100
0
10
20
30
40

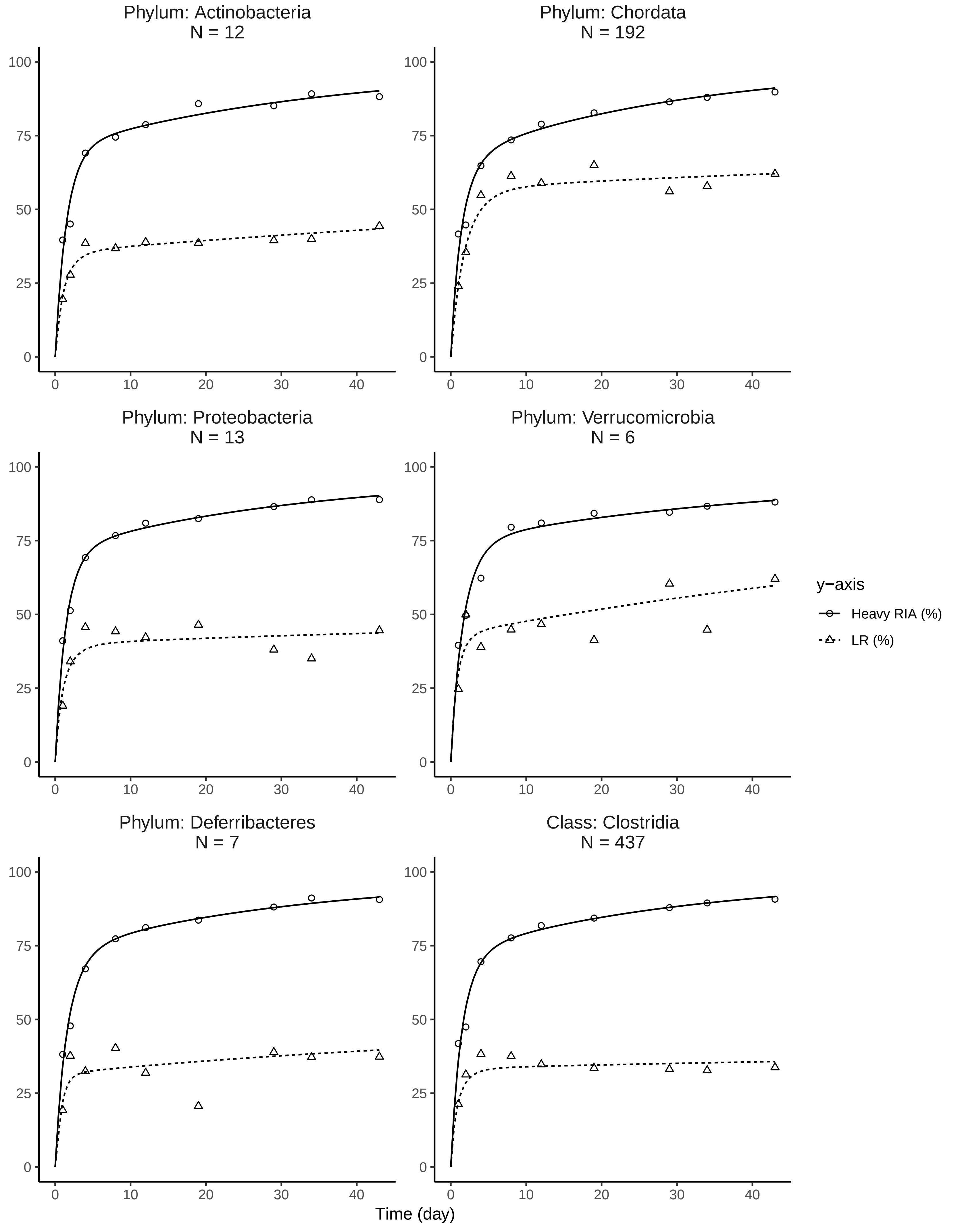

Phylum: Actinobacteria
N = 12
Phylum: Chordata
N = 192
Phylum: Proteobacteria
N = 13
Phylum: Verrucomicrobia
N = 6
Phylum: Deferribacteres
N = 7
Class: Clostridia
N = 437
0
25
50
75
100
0
10
20
30
40
Time (day)
y−axis
Heavy RIA (%)
LR (%)

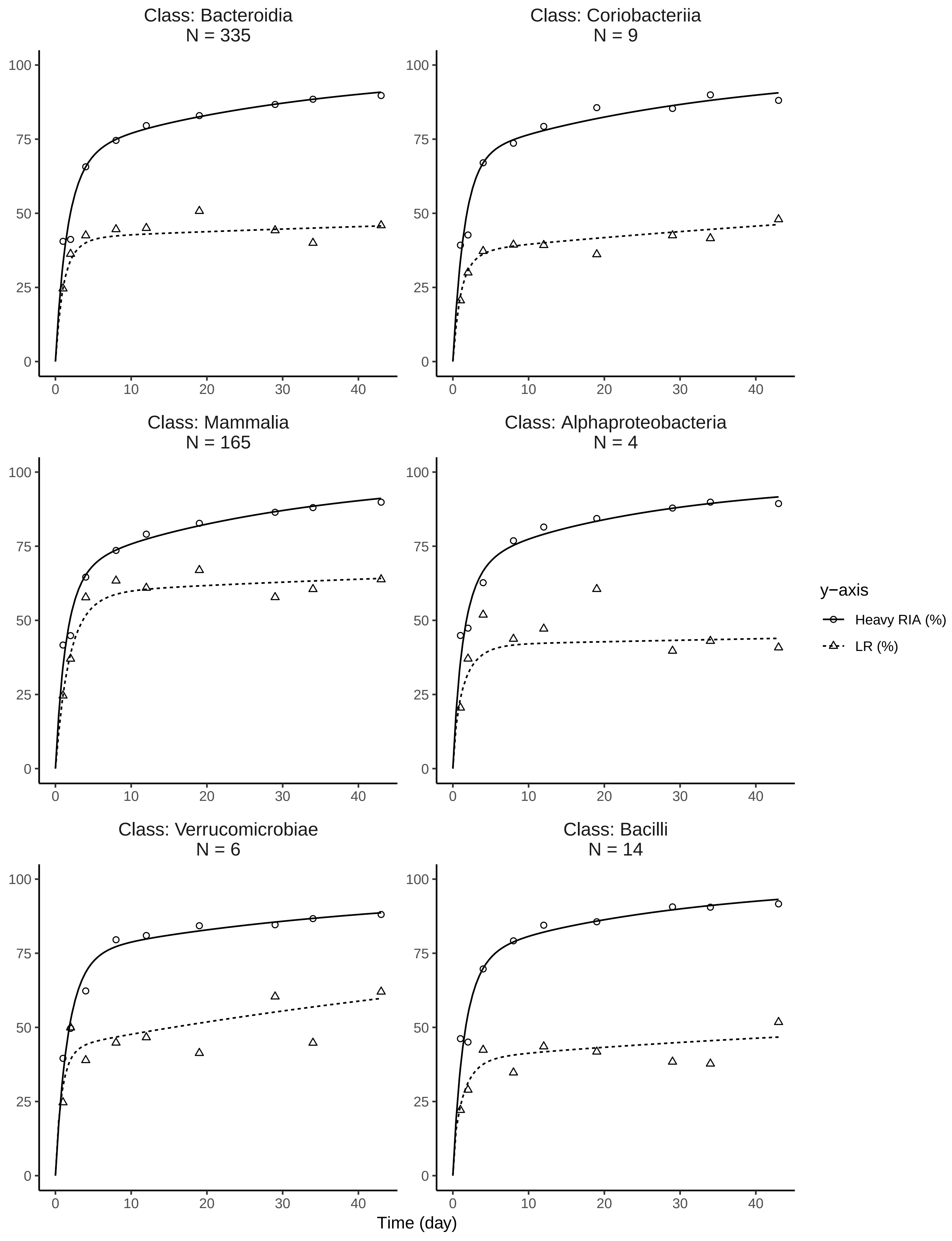

Class: Bacteroidia
N = 335
Class: Coriobacteriia
N = 9
Class: Mammalia
N = 165
Class: Alphaproteobacteria
N = 4
Class: Verrucomicrobiae
N = 6
Class: Bacilli
N = 14
y−axis
Heavy RIA (%)
LR (%)
Time (day)

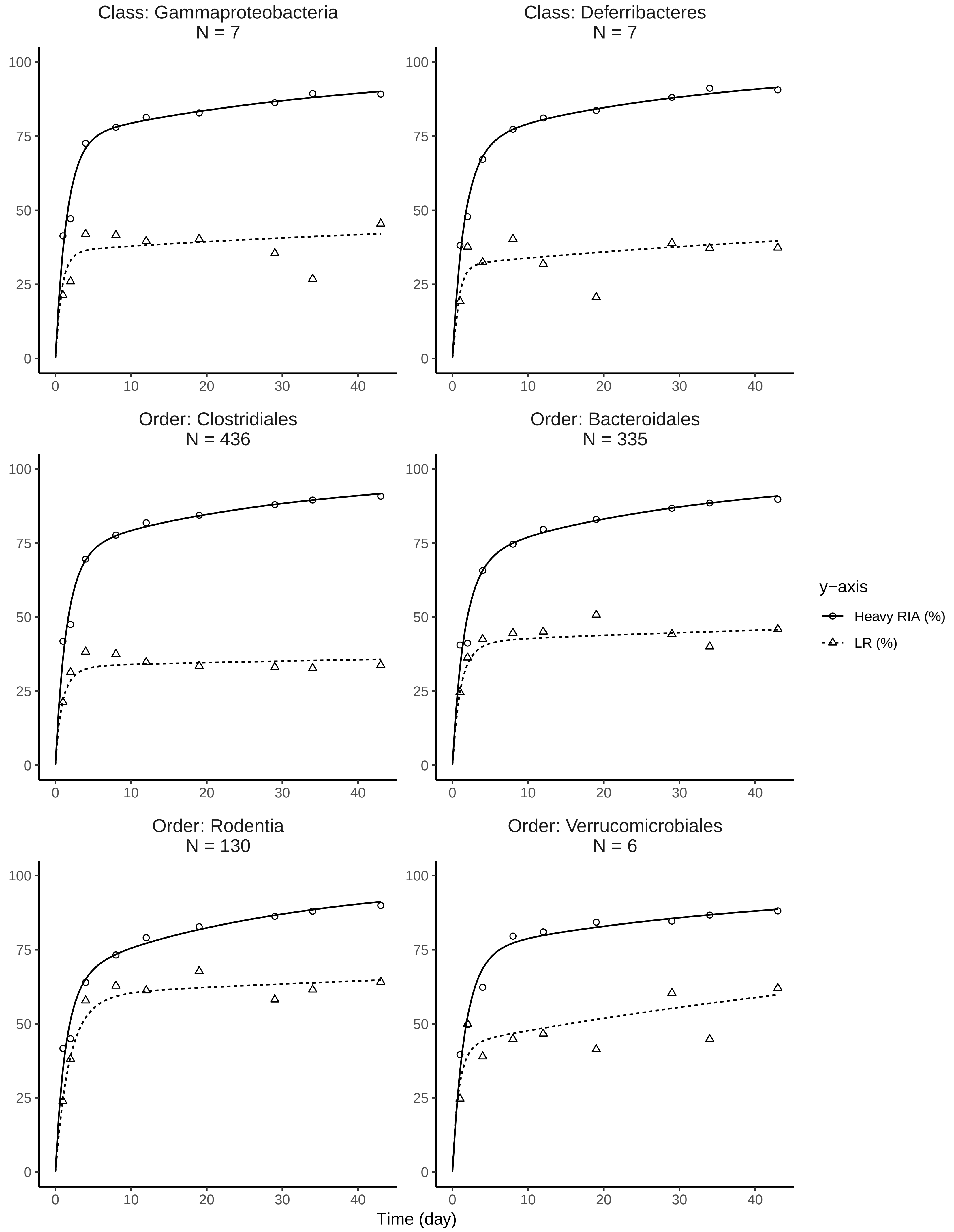

Class: Gammaproteobacteria
N = 7
Class: Deferribacteres
N = 7
Order: Clostridiales
N = 436
Order: Bacteroidales
N = 335
Order: Rodentia
N = 130
Order: Verrucomicrobiales
N = 6
y−axis
Heavy RIA (%)
LR (%)
Time (day)
0
25
50
75
100
0
10
20
30
40

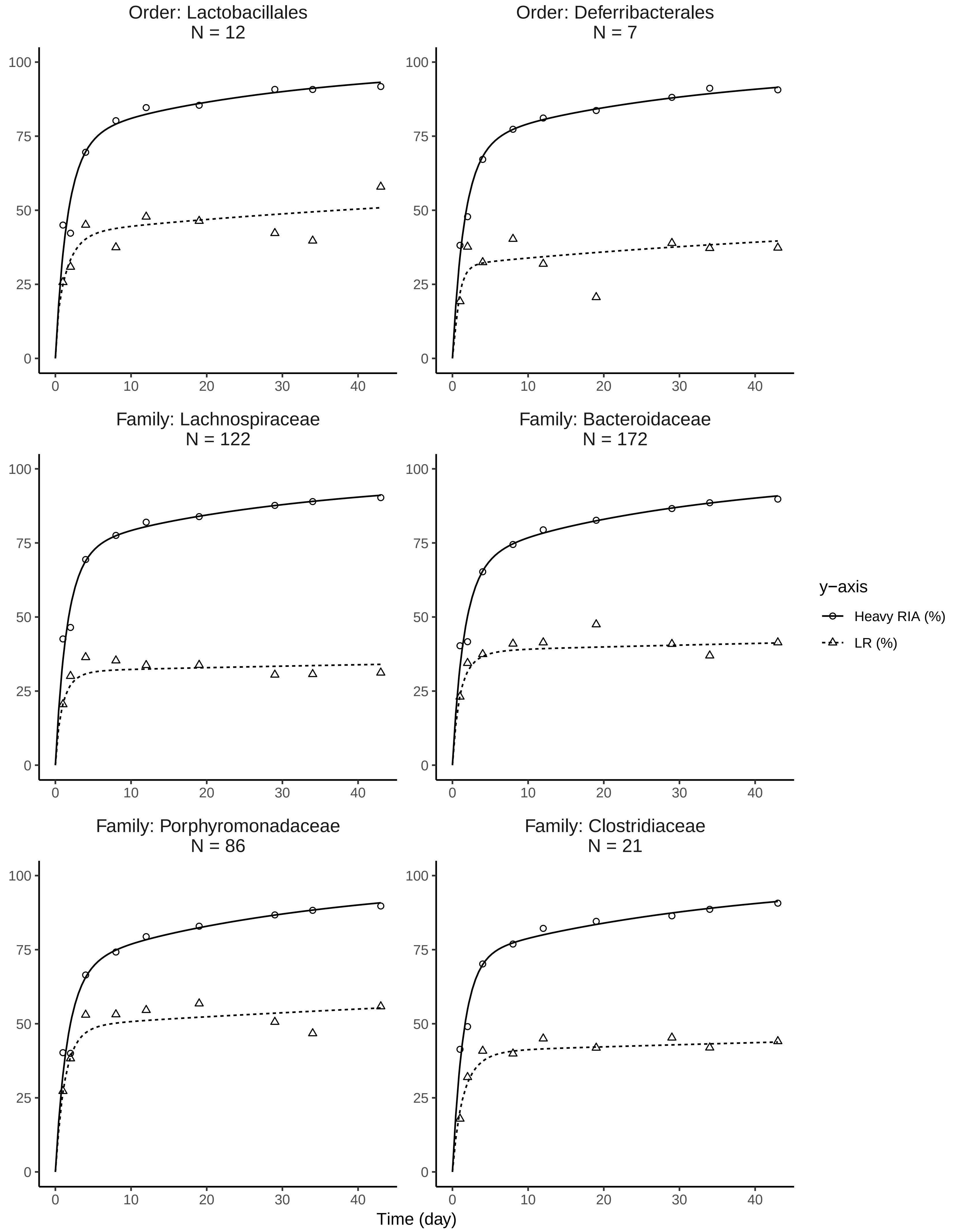

Order: Lactobacillales
N = 12
Order: Deferribacterales
N = 7
Family: Lachnospiraceae
N = 122
Family: Bacteroidaceae
N = 172
Family: Porphyromonadaceae
N = 86
Family: Clostridiaceae
N = 21
y−axis
Heavy RIA (%)
LR (%)
0
25
50
75
100
0
10
20
30
40
Time (day)

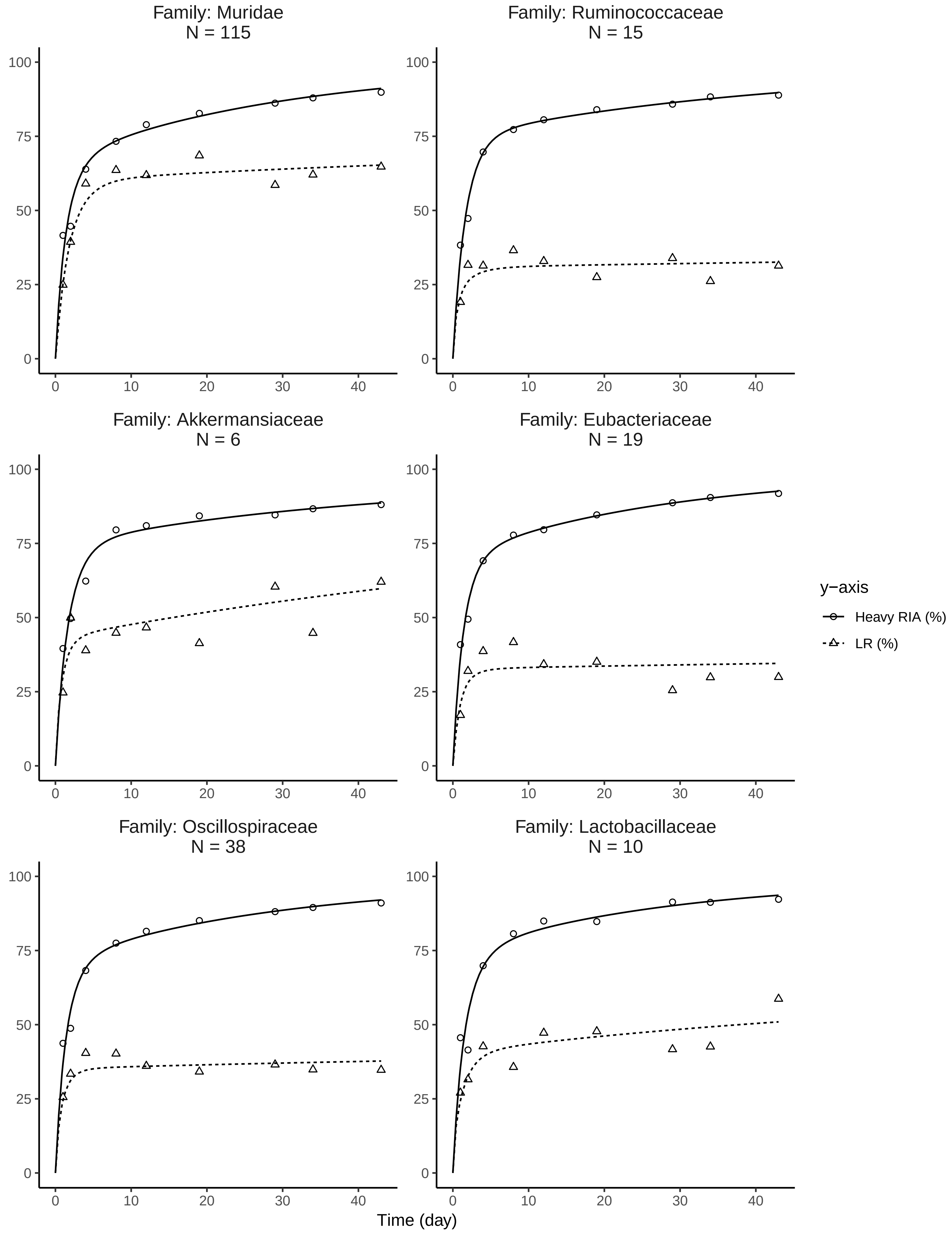

Family: Muridae
N = 115
Family: Ruminococcaceae
N = 15
Family: Akkermansiaceae
N = 6
Family: Eubacteriaceae
N = 19
Family: Oscillospiraceae
N = 38
Family: Lactobacillaceae
N = 10
0
25
50
75
100
0
10
20
30
40
y−axis
Heavy RIA (%)
LR (%)
Time (day)

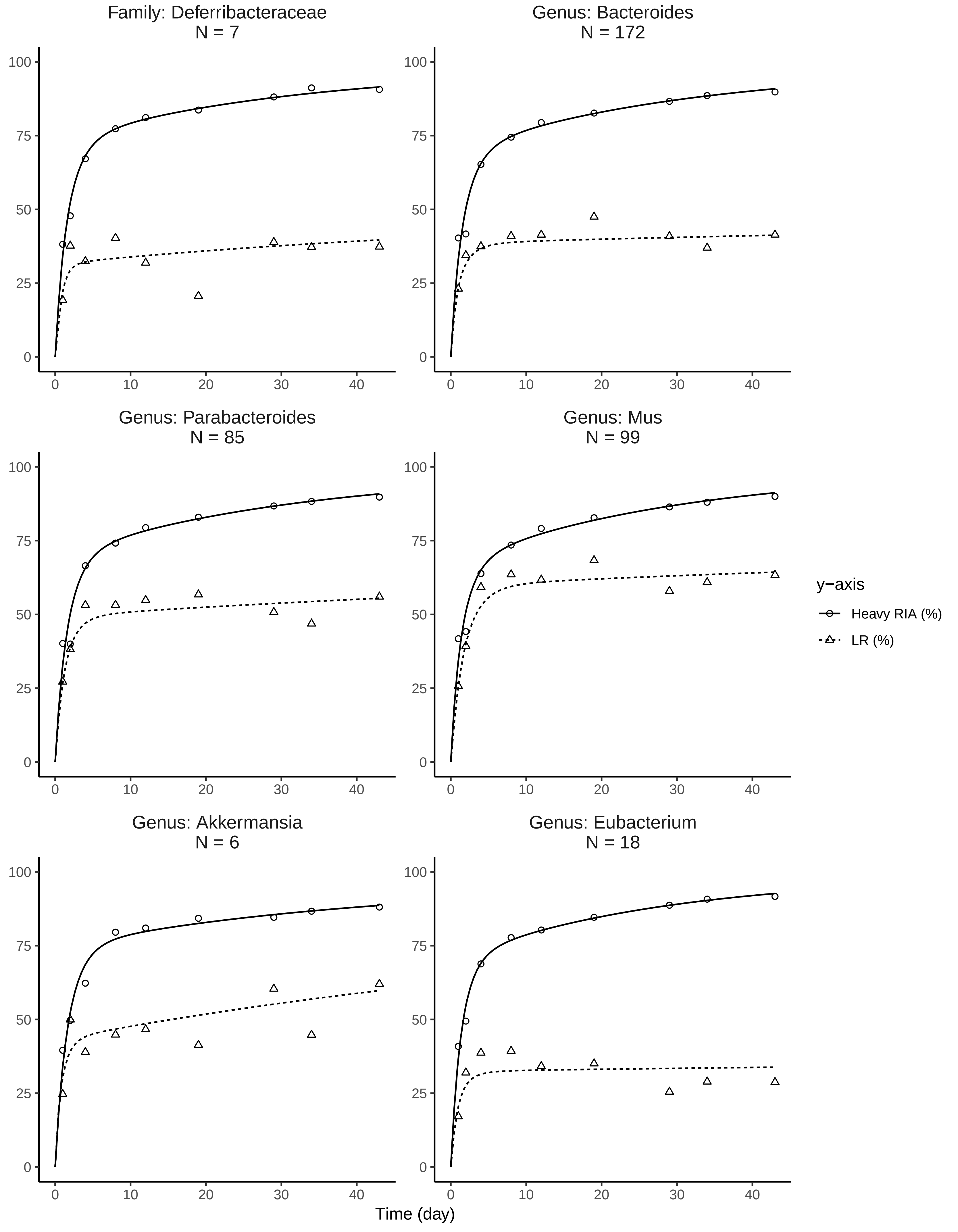
Family: Deferribacteraceae
N = 7
Genus: Bacteroides
N = 172
Genus: Parabacteroides
N = 85
Genus: Mus
N = 99
Genus: Akkermansia
N = 6
Genus: Eubacterium
N = 18
y−axis
Heavy RIA (%)
LR (%)
Time (day)
0
25
50
75
100
0
10
20
30
40

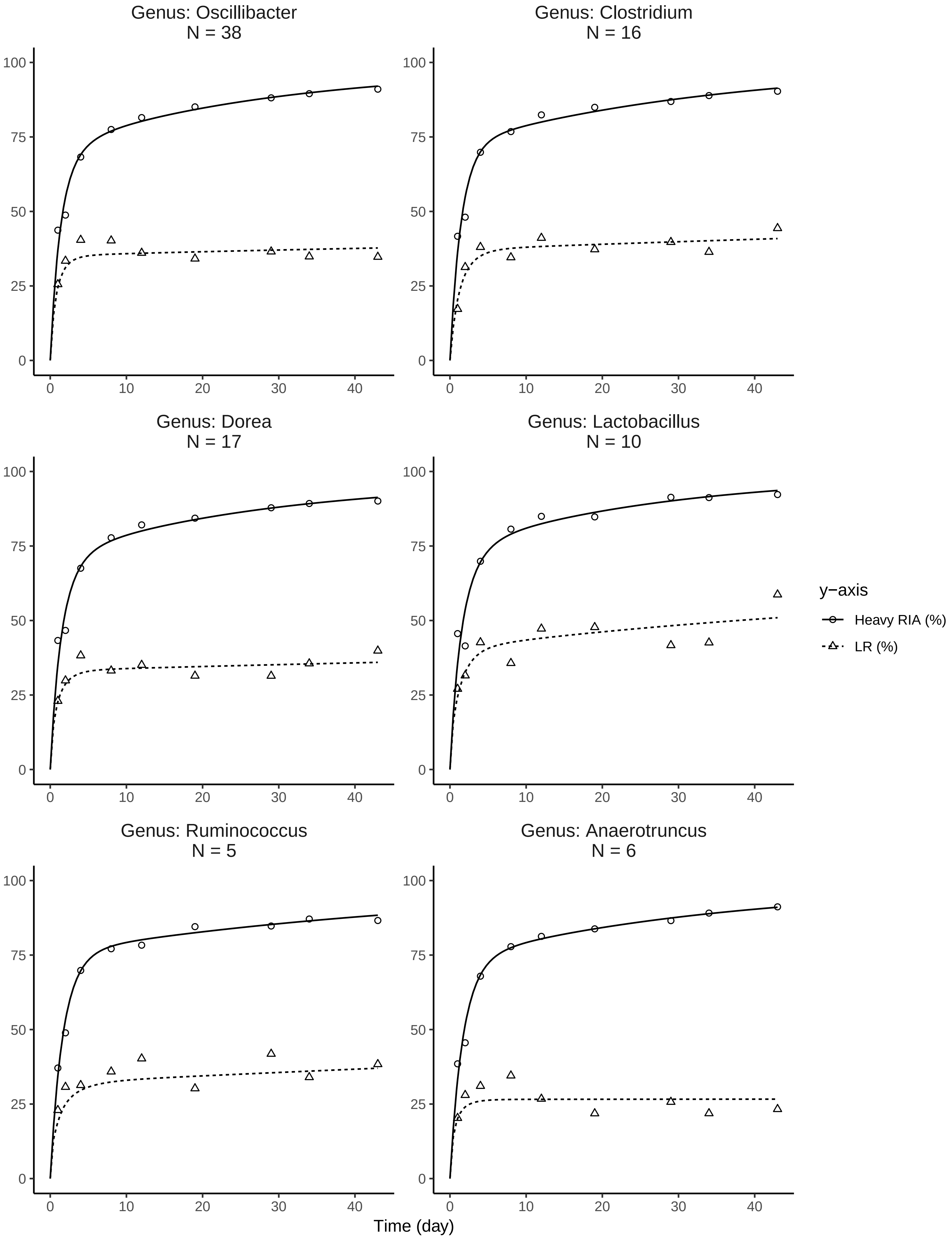

Genus: Oscillibacter
N = 38
Genus: Clostridium
N = 16
Genus: Dorea
N = 17
Genus: Lactobacillus
N = 10
Genus: Ruminococcus
N = 5
Genus: Anaerotruncus
N = 6
0
25
50
75
100
0
10
20
30
40
Time (day)
y−axis
Heavy RIA (%)
LR (%)

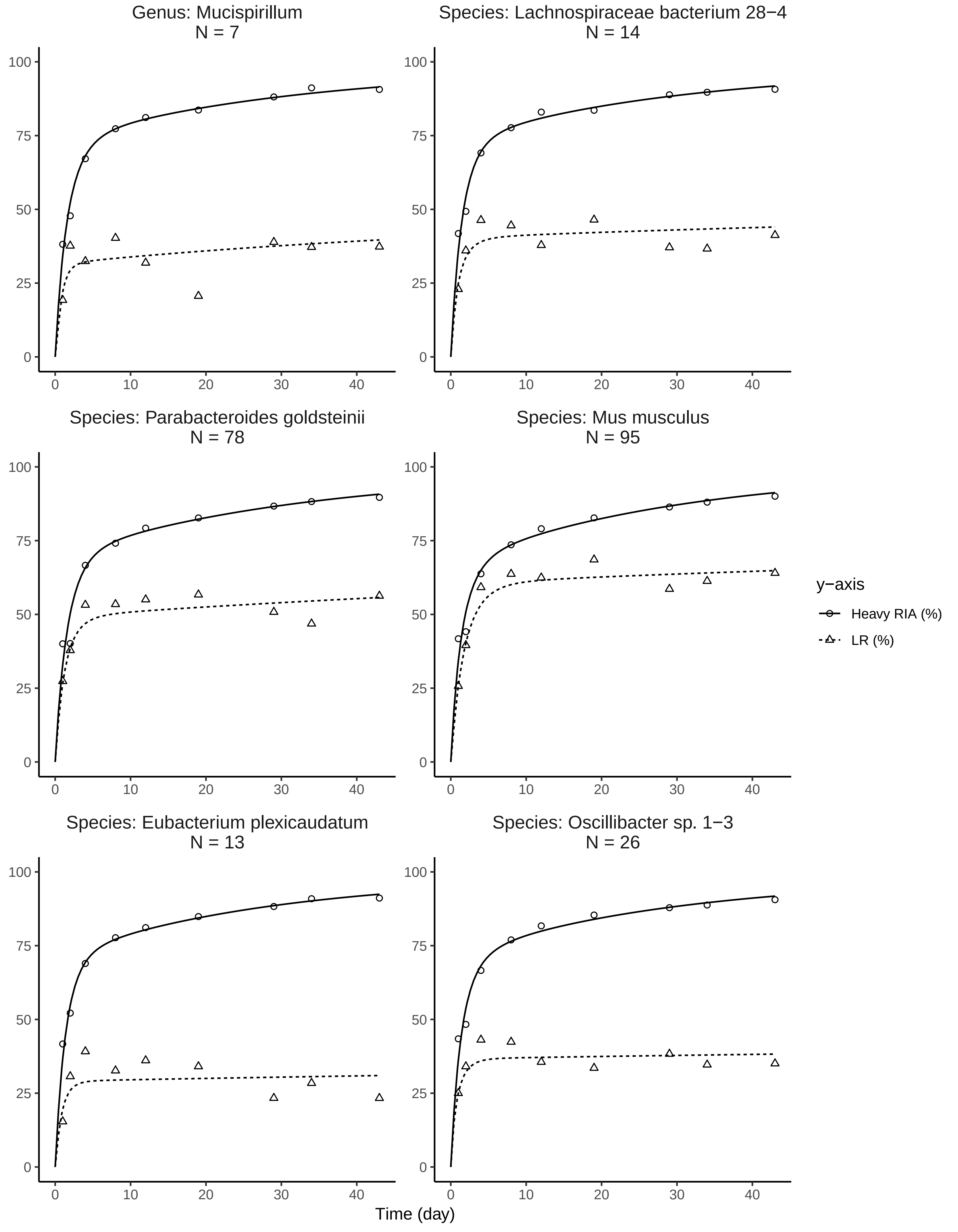
Genus: Mucispirillum
N = 7
Species: Lachnospiraceae bacterium 28−4
N = 14
Species: Parabacteroides goldsteinii
N = 78
Species: Mus musculus
N = 95
Species: Eubacterium plexicaudatum
N = 13
Species: Oscillibacter sp. 1−3
N = 26
y−axis
Heavy RIA (%)
LR (%)
Time (day)
0
10
20
30
40
0
25
50
75
100

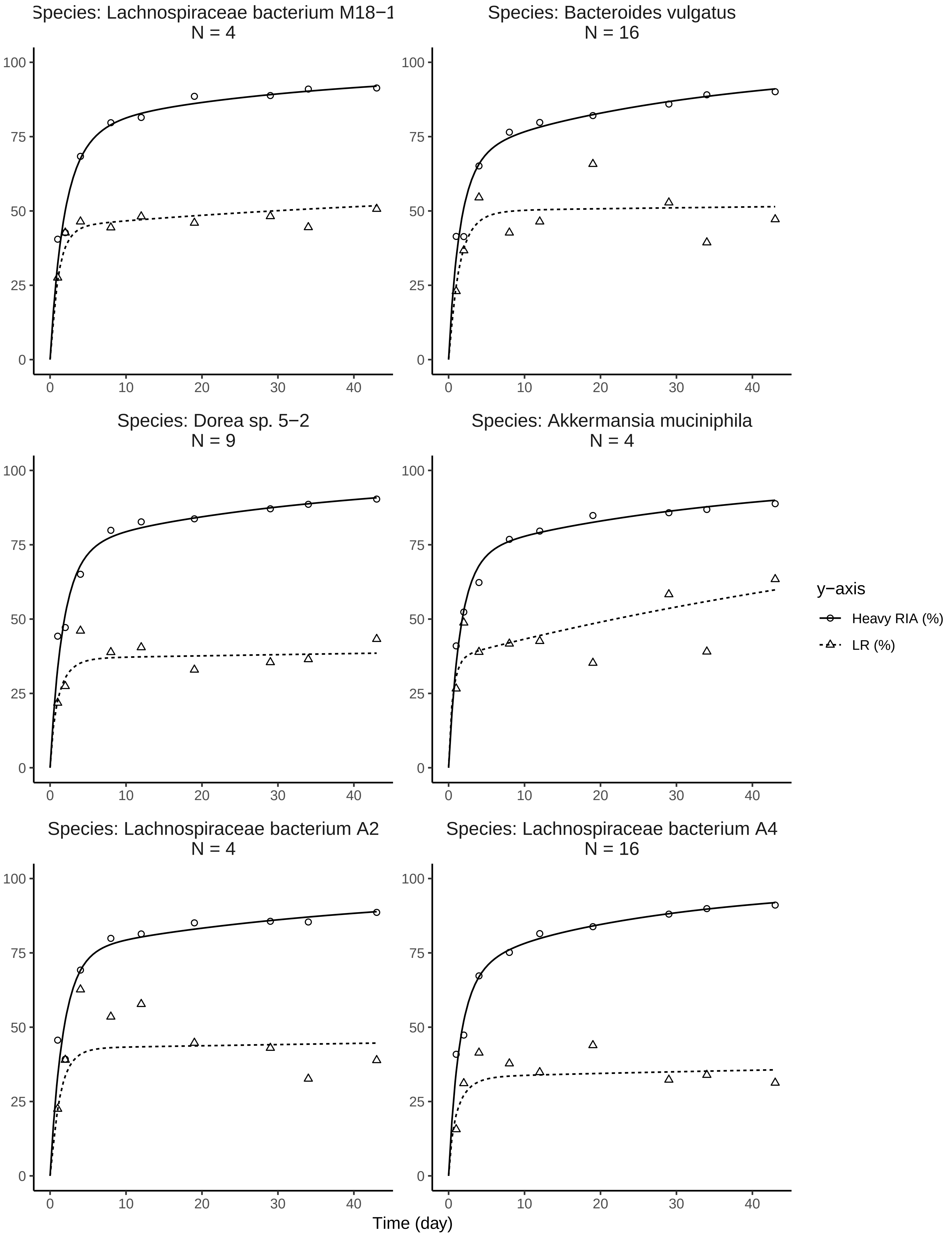

Species: Lachnospiraceae bacterium M18−1
N = 4
Species: Bacteroides vulgatus
N = 16
Species: Dorea sp. 5−2
N = 9
Species: Akkermansia muciniphila
N = 4
Species: Lachnospiraceae bacterium A2
N = 4
Species: Lachnospiraceae bacterium A4
N = 16
y−axis
Heavy RIA (%)
LR (%)
Time (day)

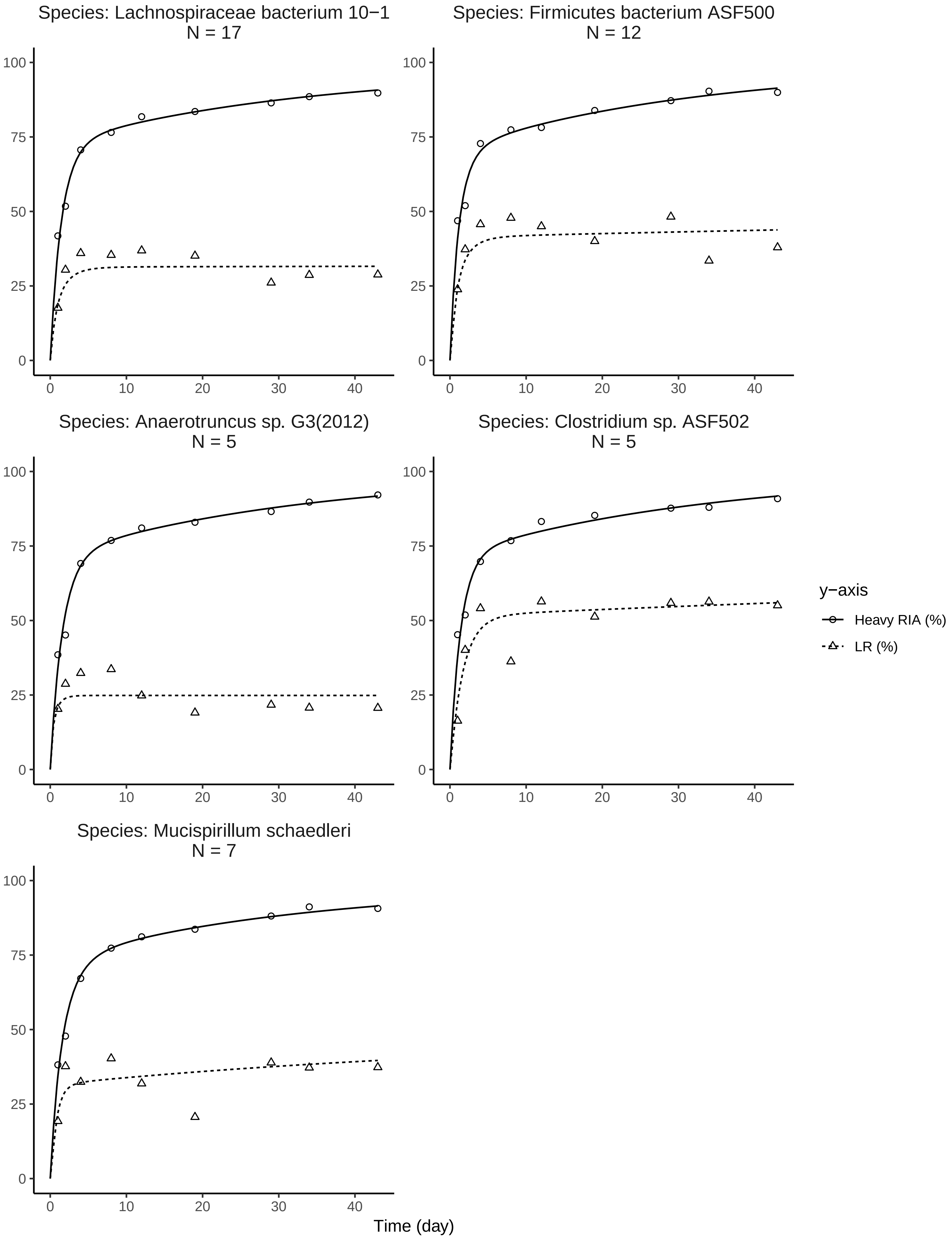
Species: Lachnospiraceae bacterium 10−1
N = 17
Species: Firmicutes bacterium ASF500
N = 12
Species: Anaerotruncus sp. G3(2012)
N = 5
Species: Clostridium sp. ASF502
N = 5
Species: Mucispirillum schaedleri
N = 7
y−axis
Heavy RIA (%)
LR (%)
Time (day)
