## Supplementary Information for "Studying the dynamics of the gut microbiota using metabolically stable isotopic labeling and metaproteomics"

### Calculating Local False Discovery Rate

Let us suppose that, at time $t$, $p_{0_t}$ is the probability of data, say $RIA_t$, with value $x_t$ for false discoveries (the null hypothesis) and $p_{1_t}$ is the probability of $RIA_t$ with value $x_t$ for true discoveries (the alternative hypothesis). In addition, let us suppose that $f_{0_t}(x_t)$ is the density of $x_t$ for the null hypothesis and $f_{1_t}(x_t)$ is the density of $x_t$ for the alternative hypothesis at time $t$. Thus, according the law of total probability, the mixture density for both hypotheses, $f(x_t)$, is measured using equation (1)

$$f(x_t) = p_{0_t} f_{0_t}(x_t) + p_{1_t} f_{1_t}(x_t). \tag{1}$$

By applying bayes rule to equation (1), we can obtain the *a posteriori* probability that the hypothesis $H$ for the random variable $RIA_t$ with value $x_t$ at time $t$ is null $p_{0_t}(x_t)$ or non-null $p_{1_t}(x_t)$.

$$\begin{aligned} p_{0_t}(x_t) &= P(H = \text{null}|RIA_t = x_t) = p_{0_t}\frac{f_{0_t}(x_t)}{f(x_t)}, \\ p_{1_t}(x_t) &= P(H = \text{non-null}|RIA_t = x_t) = p_{1_t}\frac{f_{1_t}(x_t)}{f(x_t)} = 1 - p_{0_t}\frac{f_{0_t}(x_t)}{f(x_t)}. \end{aligned} \tag{2}$$

Efron et al specify that, by definition, the LFDR, $P(H = \text{null}|RIA_t = x_t)$, is equal to $p_{0_t}(x_t)$. Thus, in order to measure LFDR, $p_{0_t}$ and $f_{0_t}(x_t)$ or $p_{1_t}$, and $f_{0_t}(x_t)$ must be measured empirically. Since the parameters of the pipeline for quantifying heavy peptide features were the same for each timepoint and that all of the heavy peptide features quantified at time zero are false, the density of RIA for false discoveries can be estimated from time zero using kernel density estimation (KDE) to obtain $f_{0_t}(x_t)$. Similarly, the probability of false discoveries at time $t$, $p_{0_t}$, can be estimated using the total number of heavy features identified at time zero, $n_{t=0}$, and the total number of heavy features identified at time t, $n_t$ Efron et al specify that, by definition, the LFDR, $P(H = \text{null}|RIA_t = x_t)$, is equal to $p_{0_t}(x_t)$. Thus, in order to measure LFDR, $p_{0_t}$ and $f_{0_t}(x_t)$ or $p_{1_t}$, and $f_{0_t}(x_t)$ must be measured empirically. Since the parameters of the pipeline for quantifying heavy peptide features were the same for each timepoint and that all of the heavy peptide features quantified at time zero are false, the density of RIA for false discoveries can be estimated from time zero using kernel density estimation (KDE) to obtain $f_{0_t}(x_t)$. Similarly, the probability of false discoveries at time $t$, $p_{0_t}$, can be estimated using the total number of heavy features identified at time zero, $n_{t=0}$, and the total number of heavy features identified at time t, $n_t$

$$p_{0_t} = \frac{n_{t=0}}{n_t}. \quad (3)$$

Finally, LFDR can be empirically measured by estimating the mixture density of value $x_t$ for false and true discoveries at time $t$ using KDE and applying $p_{0_t}$, $f_{0_t}(x_t)$, and $f_t(x)$ to equation (2). If we assume that RIA and LR are conditionally independent given the hypothesis $H$, the chain rule can be applied to obtain the joint *a posteriori* probability that the hypothesis $H$ for the random variable $RIA_t$ and $LR_t$ with value $x_t$ and $y_t$, respectively, is false, $p_{0_t}(x_t, y_t)$

$$p_{0_t}(x_t, y_t) = p_{0_t}\frac{f_{0_t}(x_t, y_t)}{f_t(x_t, y_t)} = p_{0_t}\frac{f_{0_t}(x_t)}{f_t(x_t)}\frac{f_{0_t}(y_t)}{f_t(y_t)}. \quad (4)$$

Three-Exponential Regression Model Equation

The equation derived by Guan *et al*[1], which uses a three-compartment model to find the relative fraction of $^{15}$ at each time point $\gamma(t)$, is

$$\gamma(t) = \left(1 + y_\mu e^{-\mu t} + y_v e{-}vt + y_{k_{bi}} e^{-k_{bi} t}\right) V_{max} \quad (5)$$

where

$$\begin{aligned}
\mu &= \frac{(k_{st} + k_{0a} + k_{bt}) - \sqrt{(k_{st} + k_{0a} + k_{bt})^2 - (4k_{0a}k_{bt})}}{2}, \\
v &= \frac{(k_{st} + k_{0a} + k_{bt}) + \sqrt{(k_{st} + k_{0a} + k_{bt})^2 - (4k_{0a}k_{bt})}}{2}, \\
y_\mu &= \frac{k_{0a}k_{bi}(\mu - k_{bt})}{(\mu - v)(\mu - k_{bi})\mu}, \\
y_v &= \frac{k_{0a}k_{bi}(v - k_{bt})}{(v - \mu)(v - k_{bi})v}, \\
y_{k_{bi}} &= \frac{k_{0a}(k_{bi} - k_{bt})}{(u - k_{bi})(v - k_{bi})}.
\end{aligned} \quad (6)$$

They describe that $k_{st}$ is the total protein synthesis rate constant, $k_{bt}$ is the total protein degradation rate constant, $k_{0a}$ is the amino acid out flow rate constant, and $k_{bi}$, is the individual protein degradation rate constant. Note that $V_{max}$, the maximum fraction that can be achieved, is added to the equation. For this study, $V_{max} = 100$ in order to convert the fraction into percentages.

**References**

1. Guan S, Price JC, Ghaemmaghami S, Prusiner SB, Burlingame AL. Compartment modeling for mammalian protein turnover studies by stable isotope metabolic labeling. Analytical chemistry. 2012;84(9):4014–4021.

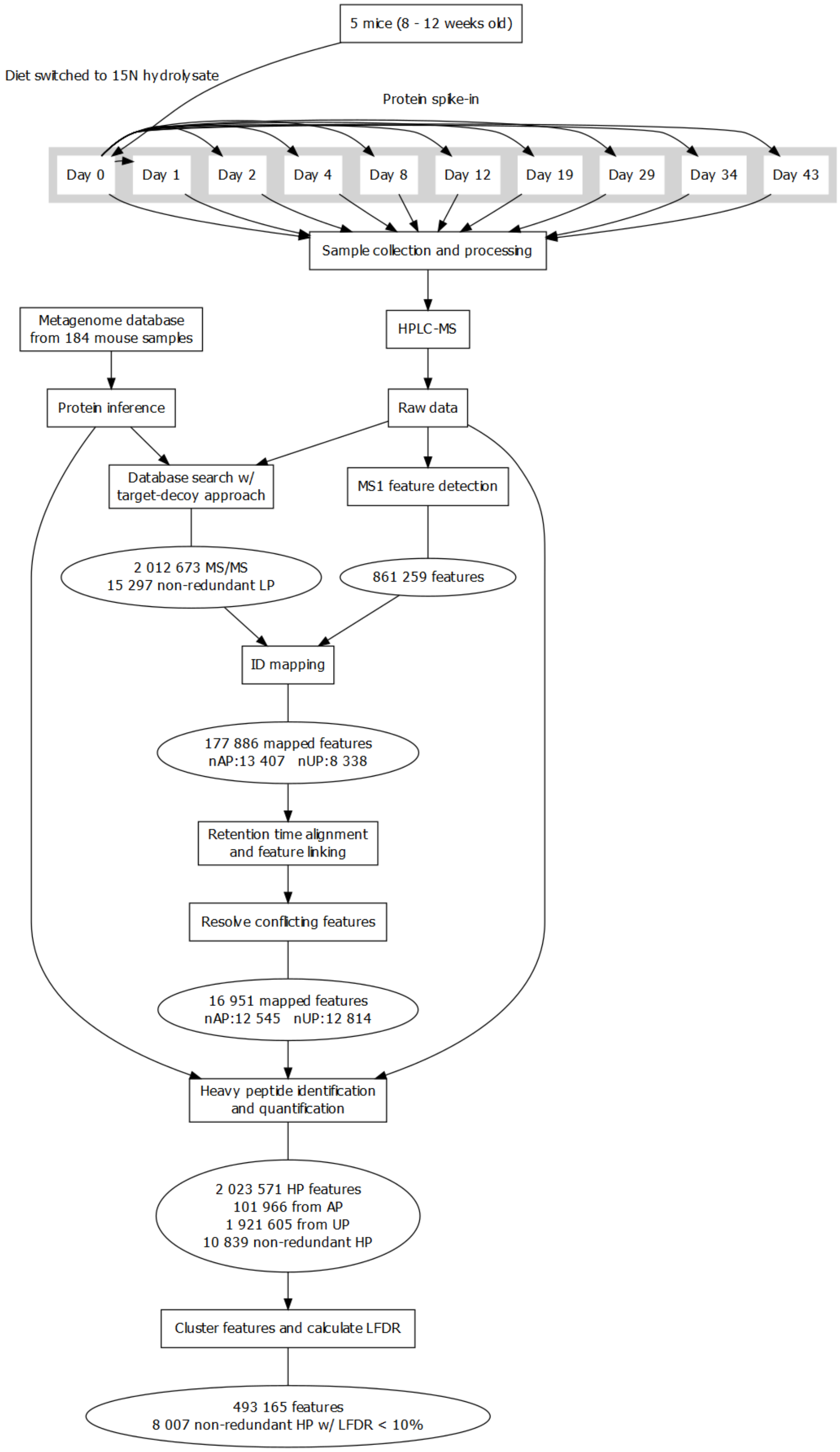


**Figure S1 Workflow Overview.** Flow chart illustrating the pipeline for identifying and quantifying heavy peptides (HP). LP = Light Peptides; AP = Assigned LP; UP = Unassigned LP; nAP = number of non-redundant LP assigned in at least one sample; nUP = number of non-redundant LP unassigned in at least one sample.

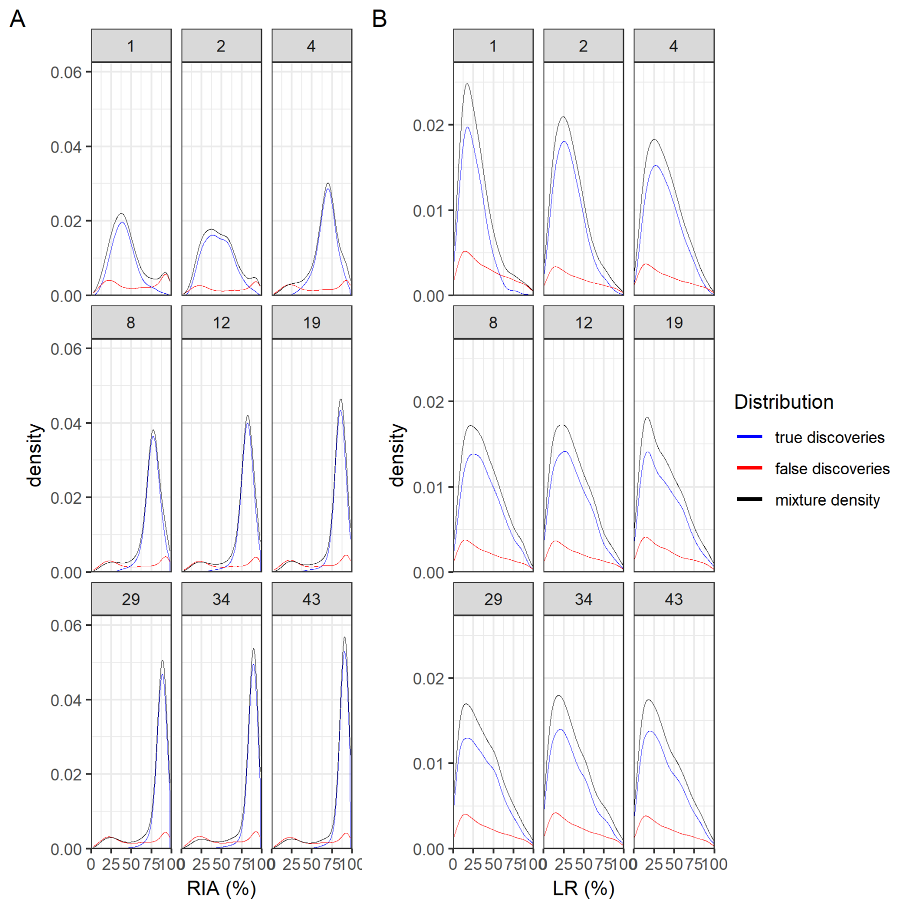


**Figure S2 LR and RIA Distribution of Mixture, False, and True discoveries.** The plot shows the density of RIA and LR for false discoveries, mixture, and estimated true discoveries.

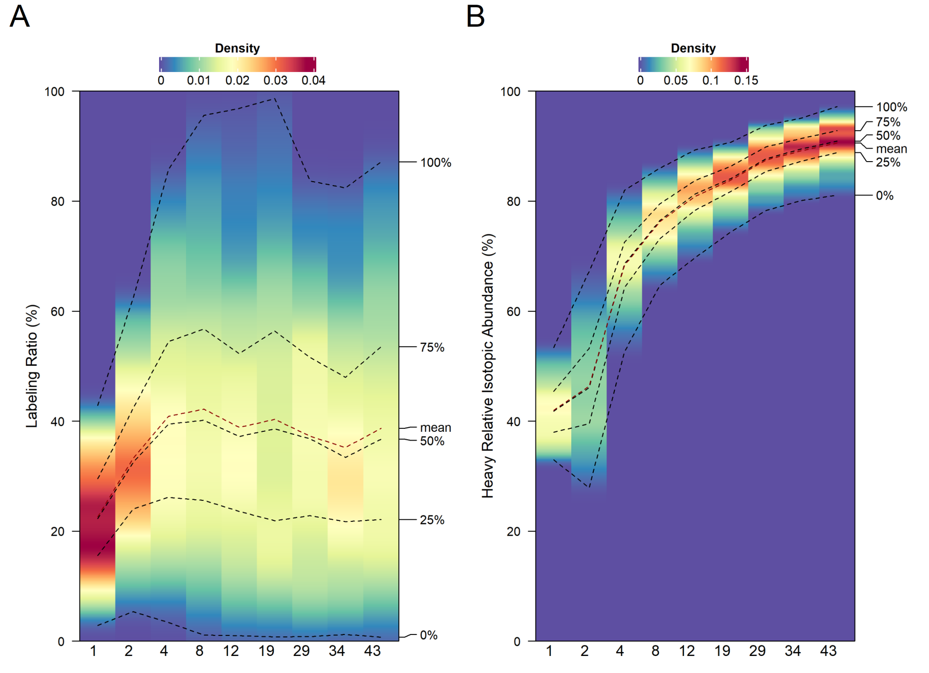


**Figure S3 RIA and LR Distribution of Heavy Peptides Below 10% LFDR Over Time.** The density distribution of A) relative isotopic distribution (RIA) and B) labeling ratio (LR) at day 1, 2, 4, 8, 12, 19, 29, 34, and 43. The black line represents the quantile of RIA and LR at 0%, 25%, 50%, 75%, and 100% and and the red line represents the mean RIA and LR over time.

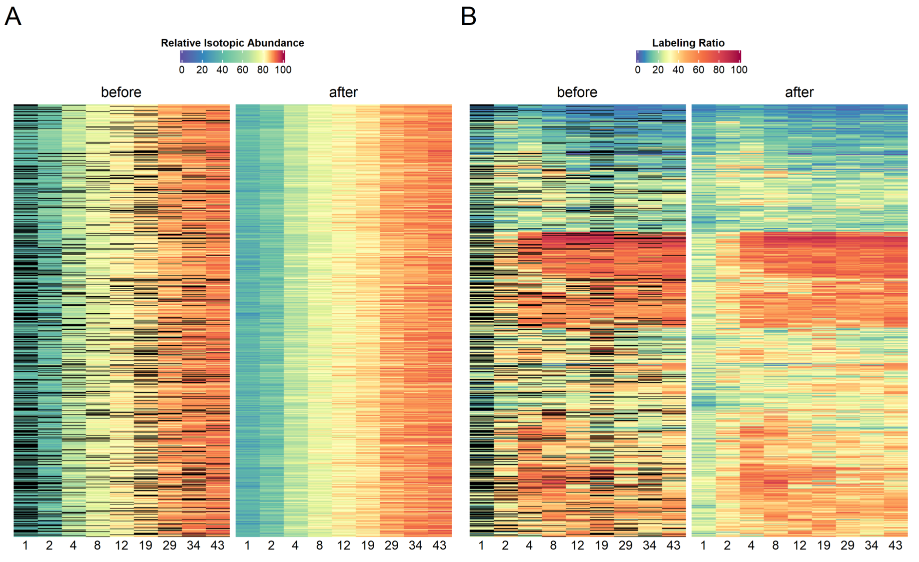


**Figure S4 Heatmap before and after imputation.** A) A heatmap denoting the distribution of relative isotopic abundance (RIA) before and after imputing using a three-exponential regression model. The columns are the timepoints and the rows are the averaged RIA of the non-redundant peptides. The color of each cell corresponds to their RIA value, as illustrated by the color key above. B) Similar to A), but with labeling ratio (LR) instead.
